# Intraspecific population admixture of a top piscivore correlates with anthropogenic alteration of freshwater ecosystems

**DOI:** 10.1101/677856

**Authors:** Erik Eschbach, Arne Wolfram Nolte, Klaus Kohlmann, Josep Alos, Sandro Schöning, Robert Arlinghaus

## Abstract

Conservation of local genetic diversity is an important policy objective, but intraspecific genetic diversity can be transformed by natural ecological processes associated with anthropogenic changes in ecosystems. Environmental changes and a strong interconnection of drainage systems impact freshwater biodiversity from gene to population level. Populations can either become extinct or expand their range and accompanying secondary contacts can lead to genetic admixture. We investigated how the genetic population structure and the patterns of genetic admixture of *Esox lucius* L. (the northern pike) vary with the type of ecosystem and the integrity of the ecosystem assessed by measures under the European Water Framework Directive. The pike inhabits river, lake and brackish water ecosystems, where it is confronted with different ecological disturbances. We analysed 1,384 pike samples from the North, Baltic and Black Sea drainages and differentiated between metapopulations from each hydrogeographic region using genotypes from 15 microsatellites and mitochondrial *cyt b* sequences. Individual populations showed signs of genetic admixture ranging from almost zero to complete replacement by foreign genotypes. Hierarchical general linear modeling revealed a highly significant positive association of the degree of genetic admixture with decreasing ecological status. This may mean that populations in disturbed environments are more prone to influences by foreign genotypes or, alternatively, increased genetic admixture may indicate adaptation to rapid environmental changes. Regardless of the underlying mechanisms, our results suggest that anthropogenic alterations of natural freshwater ecosystems can influence genetic structures, which may lead to a large-scale reduction of intraspecific genetic diversity.

## Introduction

In the age of the Anthropocene, animal and plant populations have to cope with landscapes that are used and intensively altered by humans (Christie & Knowles, 2015; Ortego et al., 2015; Sexton et al., 2013), which is discussed as the most prominent factor leading to loss of genetic diversity and evolutionary potential, and which in turn can result in the irreversible loss of populations and species (Dulvy et al., 2003; Laurance & Useche, 2009; Smith & Bernatchez, 2008). However, before extinction occurs, genetic structure of species and populations is expected to undergo changes, which may reveal processes and functions that help organisms to adapt to changing environmental conditions (Arnold, 2016).

Studies investigating fish communities and populations demonstrated that aquatic ecosystems react particularly sensitive to ecological changes (e.g. Whitehead et al., 2017), like habitat fragmentation, increasing isolation of populations through migration barriers (Waples et al., 2017), artificial opening of new routes for migration and intentional transplantation of individuals through stocking and introduction (Laikre et al., 2010). Particularly, stocking as a widespread management practice mediates direct secondary contacts among populations leading to unpredictable outcomes as a plethora of studies have shown (Allendorf et al., 2001; Diana et al., 2017; Hansen, 2002; Marie et al., 2012; van Poorten et al., 2011). According to these investigations stocked fish either disappear without a trace or become established to varying degrees, which eventually leads to admixture of non-native with native populations or even complete replacement of native populations. For example, Englbrecht et al. (2002) showed that some native populations of arctic char (*Salvelinus umbla*) were not genetically affected despite massive stocking in natural lake systems, while other natural populations were replaced by stocked char of different origin after severe eutrophication of an alpine lake. Therefore it is possible that environmental degradation through human-induced eutrophication was responsible for the natural population losing their buffering potential and rendering it vulnerable to invasion by non-native genotypes. As another example, Harbicht et al. (2014) identified a number of physico-chemical (oxygen, pH, temperature), morphometrical (surface area and depth of lakes) and topographical (elevation) factors that significantly influenced admixture in brook trout (*Salvelinus fontinalis*) as a result of stocking. Similarly, numerous studies have shown that hybridization is enhanced at the interspecific level in ecologically perturbed habitats, e.g. in African cichlids *Cichlidae* (Seehausen et al., 1997; 2008), sculpins *Cottidae* (Nolte et al., 2005), European whitefish *Corregonus spp.* (Bittner et al., 2010; Vonlanthen et al., 2012), and trout *Oncorhynchus spp.* (Heath et al., 2010).

All of these examples support the idea that the outcome of secondary contact between different populations or lineages is influenced by local ecological conditions, but as yet, little is known about their exact nature and how they might affect intra- and interspecific hybridization. Although integrative measures of ecosystem status are readily available at national and international scales and can be indicative of the environmental challenges faced by a range of taxa in the wild, research linking ecosystem status and genetic structuring of freshwater vertebrates on large geographical scales is still scarce. In Europe, the Water Framework Directive was introduced as a comprehensive policy to monitor and improve freshwater ecosystem quality and its ecological status (European Commission, 2000). Accordingly, rivers and larger lakes are regularly assessed based on a range of biological indices, from phytoplankton to fish, to assess their ecological status and inform management actions. Genetic population structures within species, however, are currently not considered in this context, although it is likely that the ecological status of water bodies is systematically related to the meta-population structure of individual species.

*Esox lucius* L., the northern pike, may well be affected by contemporary environmental change as e. g. by the loss of floodplains in rivers and nutrient inputs in lakes (Craig, 1996, Skov & Nilsson, 2018). To fulfill their life-cycle pike strongly depend on aquatic macrophytes providing shelter for early developmental stages and camouflage to hunt for prey during the juvenile stage. After reaching sexual maturation flood plains are essentially needed as spawning ground (Casselman & Lewis, 1996; Craig, 1996; Raat, 1988). It is very likely that changes in these ecological key-features affect infra- and intraspecific outcomes upon secondary contacts caused by stocking (Cowx, 1994; Guillerault et al., 2018; Hühn et al., 2014) or by active migration through canals connecting once separated water bodies (Pauwels et al., 2013). For example, Gandolfi et al. (2017) studying the invasion process of *Esox lucius* into closely related native Italian *Esox flaviae/cisalpinus* (Lucentini et al., 2011; Bianco & Delmastro, 2011) populations, observed a mosaic-type distribution of the two species and different degrees of genetic admixture, possibly as a result of the different ecological status of the studied water bodies: Lake Garda, which provides good ecological conditions for native *E. flaviae/cisalpinus* to fulfill its natural life cycle, still seems to out-compete the establishment of northern pike, while other Italian waters with a less good ecological status were strongly prone to genetic introgression. Recently in Danish pike populations at least two regional clusters were identified referring to the hydrogeographic regions of the Baltic and the North Sea, which could be further sub-divided at different river catchment scales. At the same time also deviations from native signatures were observed on local scales, prompting the authors to speculate human-assisted ecological changes affecting habitat quality as possible reasons besides historical geological alterations (Bekkevold et al. 2015; Wennerström et al., 2018).

The objective of the present study was to explore whether the ecological status of the inland waterbodies in Germany has possibly influenced the genetic structure of contemporary pike populations, particularly with respect to genetic admixture among populations of *E. lucius*. Therefore we extended the approach by Bekkevold et al. (2015) to a larger geographical scale across all principal drainage systems in Germany. More specifically, we evaluated the presence of population structure that permitted genetic assignment of individuals to their origins and used this information to identify large-scale and local signs for intra-specific admixture. Finally we tested whether the observed genetic patterns correlated with the ecological status of the water bodies assessed according to the European Water Framework Directive.

## Material & Methods

### Sampling and DNA extraction

Sampling was performed in 2011 and 2012 applying a sampling scheme that covers as many relevant water systems and types as possible over a wide geographical area in Germany, accepting that not all sampling points could be sampled with the same intensity. Nevertheless, we have established standards, i.e. a minimum of 10 individuals characterized by at least 14 microsatellites. The sample collection comprised specimens from five river catchments draining into the North Sea, six catchments draining into the Baltic Sea, and one catchment draining into the Black Sea (Table 1). Three ecosystem types were covered including 26 lakes, 24 rivers and three brackish coastal water areas (Table 1). Pike were sampled from water bodies covering the complete range of ecological states, from very good (status 1) to poor (status 5), as defined by the European Water Framework Directive (WFD, 2000/60/EC): two samples from status 1, seven from status 2, 22 from status 3, 10 from status 4 and nine samples from status 5 water bodies. 37 waters were classified as natural and 13 were classified as heavily modified (see DRYAD deposited material for details). For three small water bodies (Alte Würm, Kleiner Döllnsee, and Schulzensee) no data according to the Water Framework Directive were available because they are only generated for standing water bodies beyond 50 ha in size.

**Table 1:** Sampled water bodies, type of water body and geographic positions of sampling sites within each of the three hydro-geographic regions. Sample identification (ID) is given by a three letter code, which is used throughout the text.

| Catchment | Waterbody | Type | ID | LAT | LONG | IC |
| --- | --- | --- | --- | --- | --- | --- |
| <u>Black Sea HGR:</u> |  |  |  |  |  |  |
| Danube | Alte Würm | r | AWU | 48°13' N | 11°27' E | **BY** |
|  | Ammersee | l | <u>AMS</u> | 48°00' N | 11°07' E | **BY** |
|  | Amper | r | AMP | 48°27' N | 11°49' E | **BY** |
|  | Chiemsee | l | CHS | 47°52' N | 12°27' E | **BY** |
|  | Danube | r | DON | 48°44' N | 11°09' E | **BY** |
|  | Ilz | r | ILZ | 48°38' N | 13°26' E | **BY** |
|  | Inn | r | INN | 48°14' N | 12°59' E | **BY** |
|  | Kochelsee | l | KOS | 47°39' N | 11°21' E | **BY** |
|  | Naab | r | <u>NAB</u> | 49°05' N | 11°56' E | **BY** |
|  | Rott | r | <u>ROT</u> | 48°23' N | 12°45' E | **BY** |
|  | Starnberger See | l | STS | 47°53' N | 11°18' E | **BY** |
|  | Waginger See | l | WAS | 47°56' N | 12°46' E | **BY** |
|  | Walchen See | l | WLS | 47°35' N | 11°20' E | **BY** |
| <u>Baltic Sea HGR:</u> |  |  |  |  |  |  |
| Barthe | NN Lake (Barthe) | l | <u>BAR</u> | 54°16' N | 12°45' E | MV |
| Oder | Neiße | r | NEI1 | 51°54' N | 14°41' E | BB |
|  |  | r | <u>NEI2</u> | 51°57' N | 14°43' E | BB |
|  |  | r | <u>NEI4</u> | 52°03' N | 14°45' E | BB |
|  | Oder | r | <u>ODE2</u> | 53°03' N | 14°18' E | BB |
|  |  | r | ODE7 | 52°11' N | 14°41' E | BB |
|  | Werbellinsee | l | WBS | 52°54' N | 13°41' E | BB |
| Peene | Peene | r | <u>PEE</u> | 53°53' N | 13°28' E | MV |
| Schwentine | Großer Plöner See | l | <u>GPS</u> | 54°08' N | 10°23' E | SH |
| Trave | Großer Ratzeburger See | l | <u>GRA</u> | 53°43' N | 10°45' E | SH |
| Ucker | Hardenbecker Haussee | l | <u>HAH</u> | 53°14' N | 13°31' E | BB |
|  | Ucker | r | UCK4 | 53°31' N | 13°59' E | MV |
| - | Schaproder Bodden | c | <u>BAL2</u> | 54°30' N | 13°07' E | **D** |
| - | Schlei | c | BAL3 | 54°30' N | 09°40' E | **D** |
| - | Stettiner Haff | c | BAL4 | 53°48' N | 14°04' E | **D** |
| <u>North Sea HGR:</u> |  |  |  |  |  |  |
| Eider | Eider | r | <u>EID1</u> | 54°19' N | 09°09' E | SH |
| Elbe | Drewitzer See | l | DRS | 53°32' N | 12°21' E | MV |
|  | Elbe | r | <u>ELB6</u> | 53°12' N | 10°57' E | NI |
|  |  | r | ELB7 | 51°51' N | 12°27' E | SN |
|  | Gülper See (Havel) | l | HAV1 | 52°44' N | 12°15' E | BB |
|  | Großer Kossenblatter See | l | GKB | 52°08' N | 14°06' E | BB |
|  | Großer Stechlinsee | l | GST | 53°09' N | 13°01' E | BB |
|  | Jäglitz | r | JAG | 52°52' N | 12°24' E | BB |
| <u>North Sea HGR (cont.):</u> |  |  |  |  |  |  |
|  | Karthane | r | KAR | 52°59' N | 11°47' E | BB |
|  | Kleiner Döllnsee | l | KDO | 52°59' N | 13°34' E | BB |
|  | Krainke | r | KRK | 53°13' N | 11°04' E | NI |
|  | Müritz | l | <u>MUR</u> | 53°25' N | 12°41' E | MV |
|  | Schulzensee | l | SUS | 53°09' N | 13°15' E | BB |
|  | Schwarze Elster | r | STP | 51°28' N | 13°26' E | BB |
|  | Wittensee | l | <u>WIS</u> | 54°23' N | 09°44' E | SH |
| Ems | Ems | r | EMS3 | 52°58' N | 07°18' E | NI |
|  | Hieve | l | <u>EMS2</u> | 53°24' N | 07°16' E | NI |
| Rhine | Main | r | <u>MAI7</u> | 50°01' N | 10°31' E | BY |
|  | Rhine | r | <u>RHE2</u> | 49°09' N | 08°22' E | BW |
|  | Lake Constance | l | <u>BOS1</u> | 47°41' N | 09°02' E | BW |
|  |  | l | BOS3 | 47°43' N | 09°13' E | BW |
|  |  | l | BOS4 | 47°33' N | 09°37' E | BW |
|  |  | l | BOS5 | 47°35' N | 09°31' E | BW |
| Weser | Edersee | l | <u>EDS</u> | 51°11' N | 09°03' E | HE |
|  | Steinhuder Meer | l | <u>STM</u> | 52°28' N | 09°19' E | NI |

Type = type of waterbody: r = river, l = lake, c = coast, NN = no name; ID = sample identification code; geographic coordinates: LAT = latitude north (N), LONG = longitude east (E); IC = German federal state identification code: BB = Brandenburg, BW = Baden-Württemberg, BY = Bavaria, HE = Hesse, MV = Mecklenburg-Vorpommern, NI = Lower Saxony, SH = Schleswig-Holstein, SN = Saxony; HGR = hydro-geographic region; NN = name unknown.

Fin and muscle tissue samples of pike were collected by commercial and recreational fishers, research organizations and state fishery authorities. Samples obtained as frozen tissues were thawed in absolute ethanol (Thomas Geyer, Renningen, Germany) at room temperature and subsequently transferred to fresh ethanol following Eschbach (2012). Samples from research organizations were generally obtained preserved in ethanol, while samples from anglers were obtained air-dried. DNA of all types of samples was extracted with the nexttec^TM^ DNA isolation kit (Biozym Scientific GmbH, Hess. Oldendorf, Germany) according to the manufacturer’s instruction.

### Genetic marker analysis

We employed nuclear as well as mitochondrial markers to infer population structure and to compare our data with published data. Fifteen polymorphic microsatellites (Table 2) for pike were selected according to Eschbach & Schöning (2013). These were employed to analyze a subset of 1,384 samples of 53 populations with an average sample size of 22.1 ± 9.8 (mean ± SD) individuals per population, and 96.4 ± 94.0 (mean ± SD) individuals per river catchment. Microsatellites were co-amplified in multiplex PCR (Table 2) with a Thermocycler T Gradient machine (Biometra, Goettingen, Germany) using the Qiagen® Multiplex PCR Kit (Qiagen, Hilden, Germany). Forward primers were 5’-labeled with fluorescent dyes HEX, NED or FAM (SMB Services in Molecular Biology GmbH, Berlin, Germany) (Table 2). PCR started with 15 min at 95°C, followed by 35 cycles of 0.5 min at 94°C, 1.5 min at 58°C, 1.5 min at 72°C, and finishing with 10 min at 72°C. Fragments were sized with an Applied Biosystems 3500xL Sequencer equipped with a 24-capillary array. Chromatograms were evaluated with GeneMapper® Software v4.1 (Life Technologies, Darmstadt, Germany).

**Table 2:**
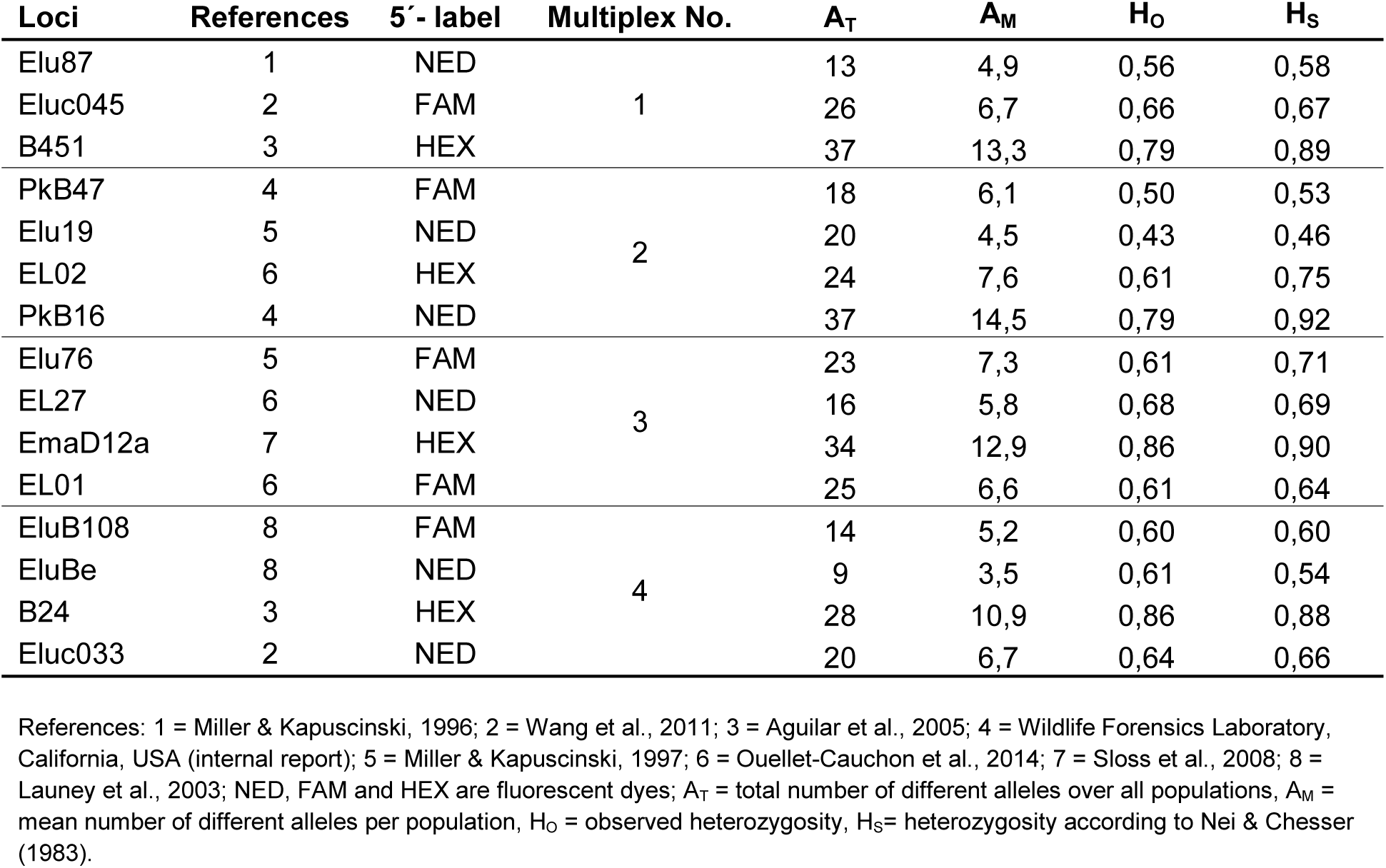
High-resolution microsatellites selected according to Eschbach & Schöning (2013) for population genetic analysis of species with low genetic variability.

| Loci | References | 5'-label | Multiplex No. | A<sub>T</sub> | A<sub>M</sub> | H<sub>O</sub> | H<sub>S</sub> |
| --- | --- | --- | --- | --- | --- | --- | --- |
| Elu87 | 1 | NED | 1 | 13 | 4,9 | 0,56 | 0,58 |
| Eluc045 | 2 | FAM |  | 26 | 6,7 | 0,66 | 0,67 |
| B451 | 3 | HEX |  | 37 | 13,3 | 0,79 | 0,89 |
| PkB47 | 4 | FAM | 2 | 18 | 6,1 | 0,50 | 0,53 |
| Elu19 | 5 | NED |  | 20 | 4,5 | 0,43 | 0,46 |
| EL02 | 6 | HEX |  | 24 | 7,6 | 0,61 | 0,75 |
| PkB16 | 4 | NED |  | 37 | 14,5 | 0,79 | 0,92 |
| Elu76 | 5 | FAM | 3 | 23 | 7,3 | 0,61 | 0,71 |
| EL27 | 6 | NED |  | 16 | 5,8 | 0,68 | 0,69 |
| EmaD12a | 7 | HEX |  | 34 | 12,9 | 0,86 | 0,90 |
| EL01 | 6 | FAM |  | 25 | 6,6 | 0,61 | 0,64 |
| EluB108 | 8 | FAM | 4 | 14 | 5,2 | 0,60 | 0,60 |
| EluBe | 8 | NED |  | 9 | 3,5 | 0,61 | 0,54 |
| B24 | 3 | HEX |  | 28 | 10,9 | 0,86 | 0,88 |
| Eluc033 | 2 | NED |  | 20 | 6,7 | 0,64 | 0,66 |

References: 1 = Miller & Kapuscinski, 1996; 2 = Wang et al., 2011; 3 = Aguilar et al., 2005; 4 = Wildlife Forensics Laboratory, California, USA (internal report); 5 = Miller & Kapuscinski, 1997; 6 = Ouellet-Cauchon et al., 2014; 7 = Sloss et al., 2008; 8 = Launey et al., 2003; NED, FAM and HEX are fluorescent dyes; A<sub>T</sub> = total number of different alleles over all populations, A<sub>M</sub> = mean number of different alleles per population, H<sub>O</sub> = observed heterozygosity, H<sub>S</sub> = heterozygosity according to Nei & Chesser (1983).

Haplotype analysis of the mitochondrial cytochrome b gene (*cyt b*) was carried out to link the present data set with the broad scale phylogeographic analysis by Skog et al. (2014) using the primers described by Grande *et al*. (2004). DNA was extracted with the ArchivePure DNA Cell/Tissue Kit (5 Prime GmbH, Hilden, Germany). The Multiplex PCR Kit (Qiagen GmbH, Hilden, Germany) was used for PCR and sequencing was performed with the BigDye Terminator v.3.1. Cycle sequencing Kit by Applied Biosystems^TM^, following the instructions of the manufacturers. Sequencing was carried out on an Applied Biosystems 3100x Genetic Analyzer. A 1.2 kbp region was amplified for a subset of 184 pike individuals belonging to 22 populations of 12 river catchments. The average sample size was 9.1 ± 2.1 (mean ± SD) individuals per population and 16.7 ± 8.5 (mean ± SD) per river catchment. Individual forward and reverse sequences were assembled using Seqman (DNA star package) and the resulting contigs were checked by eye to correct sequencing errors. Sequences of all main and sub haplotypes were deposited at the NCBI database (Acc. no. KY399416 – KY399442).

### Genetic data analysis

Microsatellite data were tested for the presence of null alleles with MICROCHECKER 2.2.3 (van Oosterhout et al., 2004) using 1,000 randomizations and applying a 95% confidence interval. Total and mean numbers of alleles as well as heterozygosity (H_O_ and Neís H_S_) were calculated with FSTAT 2.9.3.2 (Goudet, 1995). GENEPOP 4.2 (Raymond & Rousset, 1995) was applied to test for Hardy-Weinberg deviations and linkage disequilibrium setting the Markov chain parameters (MCMC) to 10,000 dememorizations, 20 batches and 5,000 iterations per batch.

STRUCTURE 2.3.2 (Falush et al., 2003) was used to infer the most likely population structure based on microsatellite data of 53 pike populations. The calculation was done with an admixture model without *a priori* population information, using a burn-in period of 100,000 repeats, 100,000 subsequent MCMC repeats and 10 iterations for each k value between one and 30. The most likely number of ancestral populations was identified as the k value, where the change of likelihood dropped considerably compared to subsequent values (Δk criterion). All individuals were assigned to each of the ancestral gene pools as defined in the most likely STRUCTURE model. NA describes the fraction of the genome inherited from the drainage basin-specific lineage, as opposed to ancestry that most likely originated from a different river basin according to the STRUCTURE model. To express all foreign genetic influences (irrespective of their origins) in relation to NA, hybrid indices (HI) for each individual were inferred from the individual NA values. Using the formula HI = 1 – (2 × |0.5 – NA|) results in a value of 1.0, if the native and foreign ancestries contributed equally to an individual’s genetic composition (maximal hybrid status, as found in a first generation hybrid), and a value of 0, if only the native ancestry contributed to an individual’s genetic composition.

MSA 4.05 (Dieringer & Schlötterer, 2003) was employed to calculate genetic differences (Nei’s D_A_ (Nei, 1983), chord distances (Cavalli-Sforza & Edwards, 1967) and the proportion of proportiond alleles (Bowcock et al., 1994) among all populations. To allow for bootstrapping, the permutation option was set to 10,000. Consensus trees were calculated subsequently with the NEIGHBOR and CONSENSUS packages of PHYLIP 3.695 (Felsenstein, 1981) and displayed with the software MEGA 5 (Tamura *et al*., 2011).

Principal coordinate analysis as an alternative to identify genetic clusters was performed with GeneAlEx 6.5 (Peakall & Smouse, 2012) using the covariance matrix obtained from Fst values (Table S2).

CLUSTAL X Version 2 (Larkin et al., 2007) was used to align all *cyt b* sequences along with 24 reference sequences of haplotypes described by Skog et al. (2014). The alignment was trimmed to a length of 1,174 bp that contained the sites that were diagnostic for the groups of haplotypes described by Skog et al. (2014). This alignment was used to confirm the presence or absence of the respective haplotypes in the populations studied here. The relationship among all haplotypes were visualized using a median-joining network as described by Bandelt et al. (1999) that was constructed using the program NETWORK 4.6.1.3 (Fluxus Technology Ltd, Suffolk, UK).

### Environmental effects on genetic structure

585 pike individuals from 24 lakes and 392 pike individuals from 23 rivers were tested in two independent hierarchical general linear models (HGLM) to infer the effect of different ecological predictors on native ancestry (NA) and hybridization index (HI). These response variables (*y*) were composed of values within the standard unit interval *y_i_* Є [1,0], where *i* designated an individual fish. Special techniques were required for linear modelling with respect to binomial errors and beta distributed random effects to incorporate features such as heteroskedasticity or skewness commonly observed in this type of data (Cribari-Netom & Zeileis, 2010). The data comprised repeated measures within individual water bodies (*n* individuals from 50 different water bodies), and therefore we considered the variance attributed to water bodies as a random effect. In addition, to account for a higher probability of natural exchange among individuals sampled in specific waterbodies within a basin, water bodies were nested within catchments. We then modelled HI and NA on a set of predictors using a linear predictor with unknown coefficients and a link function (logit). The predictors considered were: the type of water body (lake or river), its level of modification (not modified or highly modified) and its ecological status as a numerical covariate from 1 (very good) to 5 (poor) according to the European Water Framework Directive. The raw data were retrieved from the Federal Institute of Hydrology (BfG, Koblenz, Germany: http://geoportal.bafg.de/mapapps/resources/apps/had and http://geoportal.bafg.de/mapnavigator) and were deposited in DRYAD. We used the glmmADMB of the R-package, built on the open source AD Model Builder nonlinear fitting engine, to fit two HGLM models (one for NA and another one for HI) considering a beta response distribution type using the logit-link function (Fournier et al., 2012). The estimates of the fixed effects (in logit scale) as well as their standard error were estimated via the Laplace approximation. In the initial HGLMs, we included all two-level interactions among the predictors. Non-significant interactions were sequentially removed from the minimal adequate model testing the main effects.

## Results

### Assessment of genetic markers

All of the 15 microsatellite loci proved to be highly polymorphic with a total number of 9 to 37 different alleles over all pike populations and a mean number of 3.5 to 14.5 different alleles per population (Table 2). The potential presence of null alleles was detected in 1.6% of alleles over all loci and populations (Table S1). Diversity measures for observed (H_O_) and Nei’s (H_S_) heterozygosity ranged from 0.43 to 0.86 and 0.46 to 0.92, respectively, over all populations (Table 2). Populations showed deviation from Hardy-Weinberg equilibrium in 2.7 ± 1.9 loci (mean ± SD) reflected in significant heterozygote deficiencies in 2.7 ± 1.6 loci (Fig. S1). Linkage disequilibrium was detected in 3.3% of all possible loci combinations after Bonferroni correction (Fig. S2). Because departures were distributed over many loci and populations, all loci were used for population genetic analysis.

For network analysis, a 1,174 bp region of the mitochondrial *cyt b* gene containing 48 variable positions was selected. With a total of 918 informative sites in 208 sequences analyzed (excluding sites with gaps and missing data; including reference sequences), the overall information content was relatively low (3.8%) as expected for pike.

### Genetic structure of pike populations in Germany

Analysis of microsatellite data with STRUCTURE suggested a k value of either three or five as the most likely number of existing genetic clusters of pike (Fig. 1, showing the relevant k range only). However, five clusters were considered less likely based on the definition of the minimal Δk criterion. To further confirm the results obtained with STRUCTURE, we considered two additional genetic analyses based on microsatellite data. First, calculating genetic differences resulted in dendrograms (trees) with three main clusters. Although bootstrap values were mostly low (and therefore omitted from fig. 2), the overall topology of the consensus trees proved to be stable. Using Nei’s D_A_ yielded exactly the genetic structure of pike populations predicted by the assignments of STRUCTURE assuming three as the most likely number of k (Fig. 2). Moreover, the tree analysis showed that the severely admixed pike populations were grouped into the expected “new” genetic background, e.g. the Rhine population (RHE2) in the Baltic Sea hydrogeographic group or pike of Großer Plöner See (GPS) in the Black Sea hydrogeographic group (Fig. 2). Trees based on the two other distance measures - chord distances and proportion of shared alleles – yielded the same basic structure of trees, but showed one and three deviation/s compared to the predictions of STRUCTURE, respectively (trees not shown). Second, principal coordinate analysis (PCoA) based on Fst values was employed (Fig. 3). Although variation was moderate (accumulated variance explained by axes 1 and 3 = 23.2% and 22.6% by axes 1 and 2) a clear clustering into three groups representing the drainage basins of the North, Baltic and Black Sea, respectively (Table 3), was obtained, which supported a k value of three predicted by the STRUCTURE analysis. Using the PCoA, severely admixed populations were as well positioned in the genetic background predicted by the STRUCTURE analysis, providing further evidence for k = 3.

**Fig. 1:**
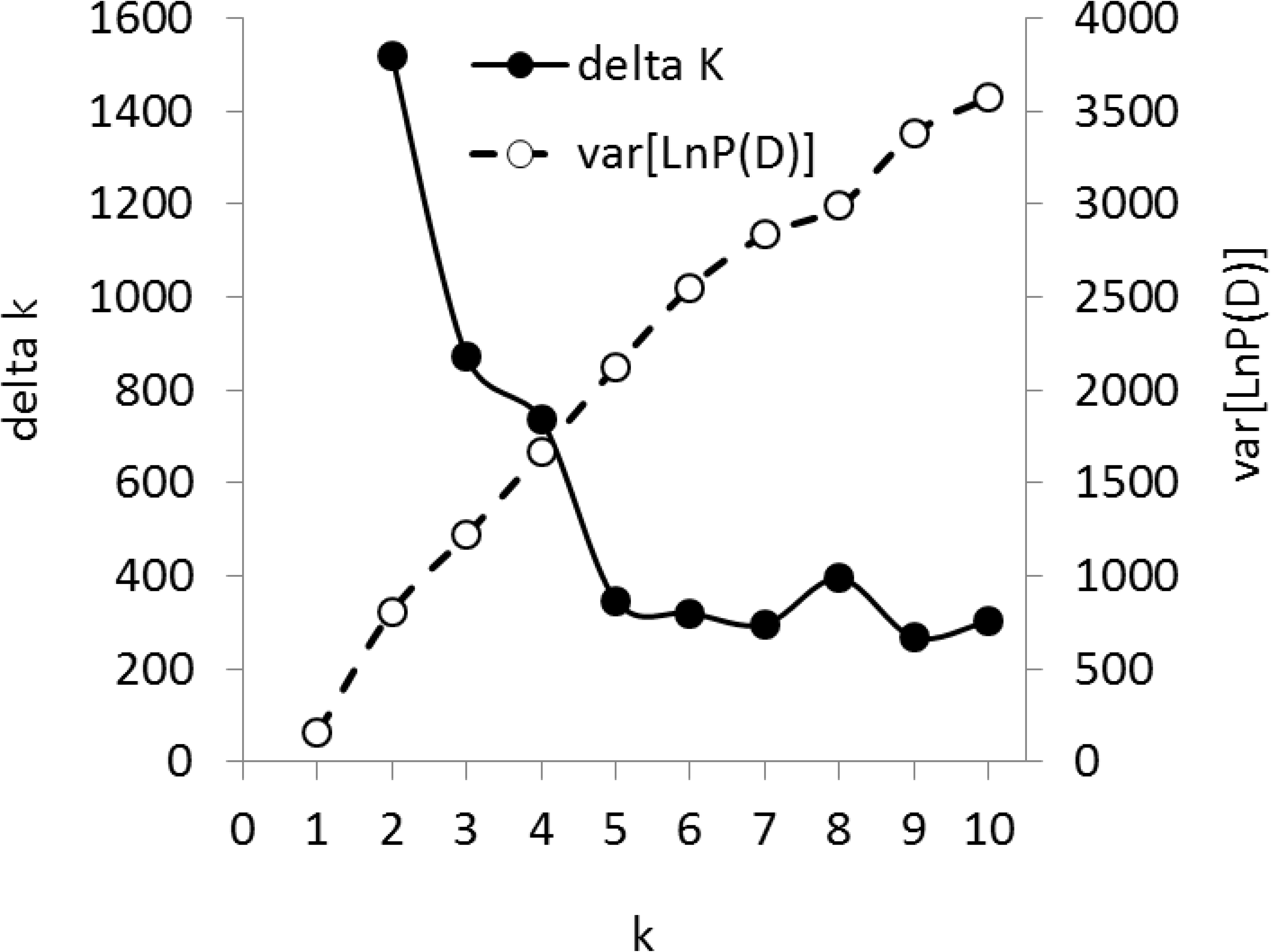
Admixture analysis revealed three or five genetic clusters as the most likely numbers, as indicated by a decrease in Δk and an increase in variance of calculated probabilities P(D). Only the relevant range of calculated k is shown here.

**Fig. 2:**
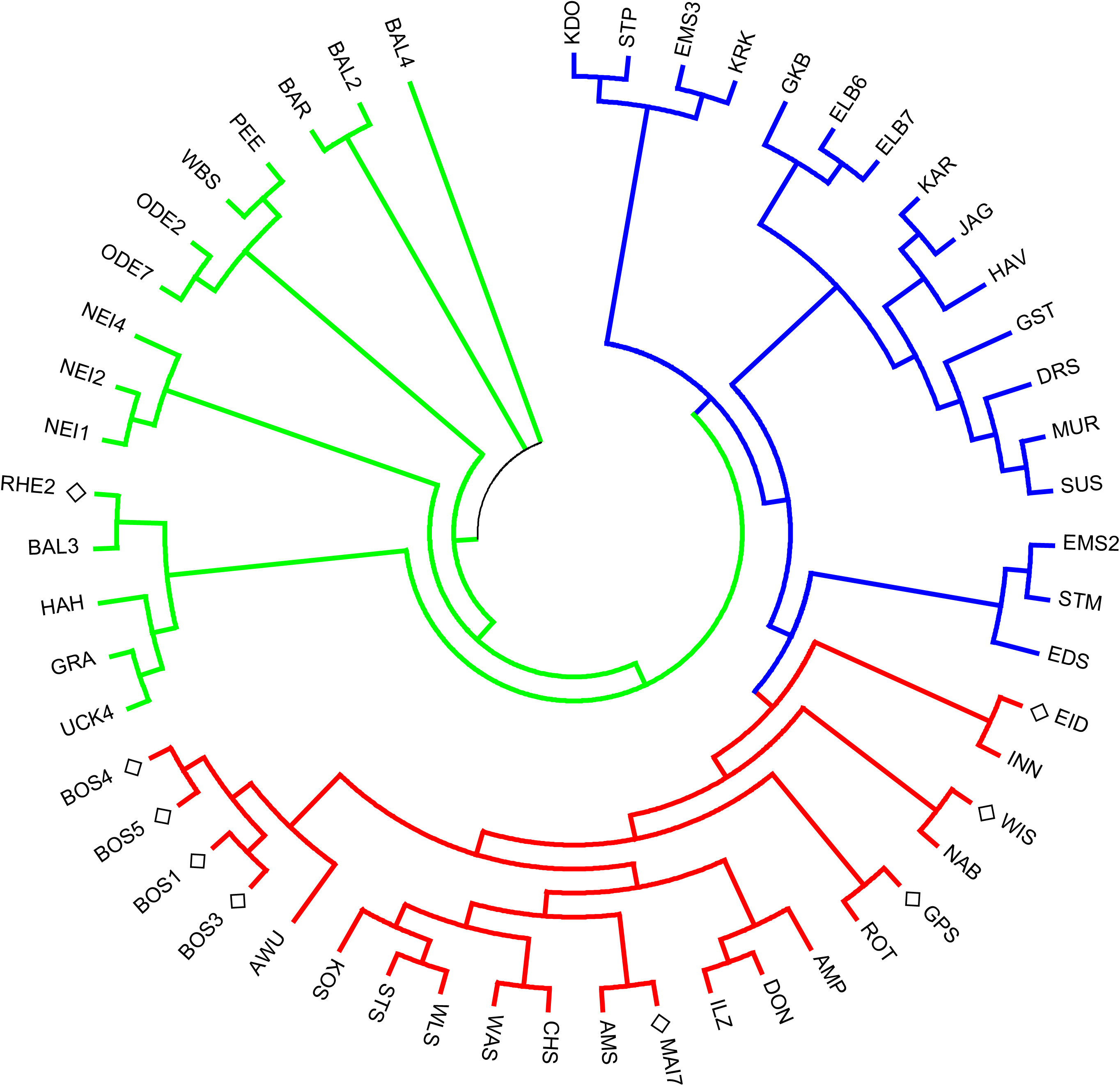
Neighbor joining consensus tree based on 10,000 permutations for calculating Neís genetic distance estimator D_A_. Although bootstrap values were generally low, the tree topology proved to be stable and was consistent with the most likely STRUCTURE model predicting three main clades. Furthermore, admixed populations (labeled with a diamond) grouped according to their predicted dominant genetic background within the respective clades, e.g. the RHE2 population of the river Rhine groups within the Baltic Sea clade (green branches) and the GPS population sampled from Großer Plöner See, which is connected with the Baltic Sea is positioned in the Black Sea clade (red branches). Branches of the North Sea clade are drawn in blue.

**Fig. 3:**
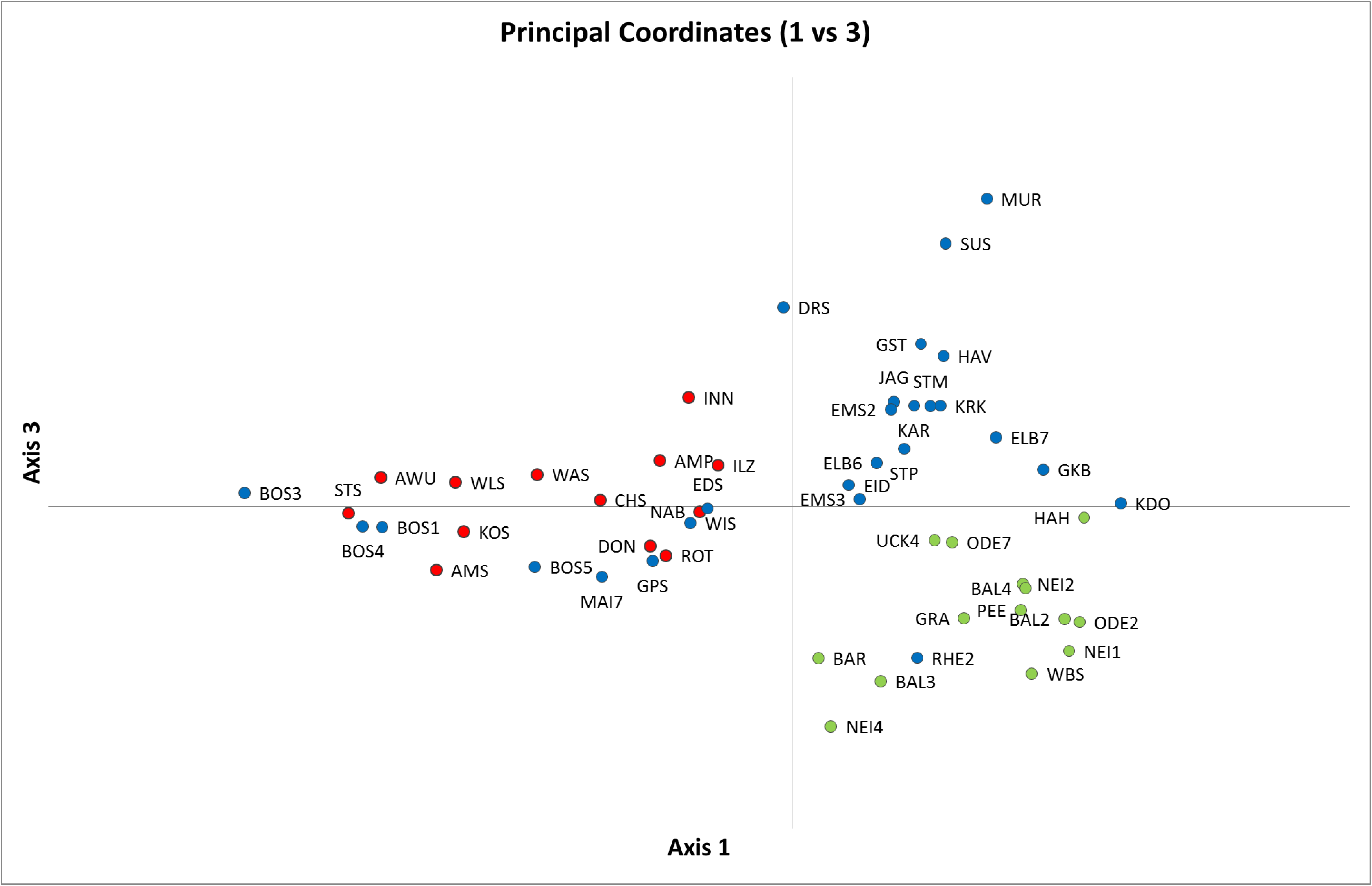
Principal coordinate analysis based on pairwise Fst values (see Table S2) of all pike samples. Despite low levels of variation (23.2% accumulated variation of axis 1 and 3) the three main clusters predicted by the most likely model of STRUCTURE were clearly resolved and admixed pike populations were positioned within the correct genetic context.

**Table 3:** Admixture analysis revealed three genetic clusters of pike populations belonging to the hydro-geographic region of the North, Baltic and Black Sea, respectively (shaded areas indicate highest proportion of ancestry). Some populations exhibited high proportions of non-native ancestry (indicated in fat italic writing). Sample IDs are explained in Table 1. Samples with the same number have been pooled for a clearer presentation in Fig. 4.

| ID* | no. in Fig. 4 | Proportion of ancestry: |  |  | ID | no. in Fig. 4 | Proportion of ancestry: |  |  |
| --- | --- | --- | --- | --- | --- | --- | --- | --- | --- |
|  |  | North Sea | Baltic Sea | Black Sea |  |  | North Sea | Baltic Sea | Black Sea |
| AMP | 1 | 0,140 | 0,048 | 0,812 | ODE2 | 3 | 0,306 | 0,598 | 0,096 |
| AWU | 1 | 0,138 | 0,040 | 0,821 | ODE7 | 3 | 0,416 | 0,487 | 0,097 |
| DON | 1 | 0,225 | 0,111 | 0,664 | NEI1 | 3 | **0,450** | 0,370 | 0,181 |
| ILZ | 1 | 0,218 | 0,108 | 0,674 | NEI2 | 3 | **0,629** | 0,180 | 0,191 |
| INN | 1 | 0,278 | 0,215 | 0,507 | NEI4 | 3 | 0,176 | 0,509 | 0,316 |
| NAB | 1 | 0,407 | 0,106 | 0,487 | WBS | 3.1 | 0,068 | 0,897 | 0,035 |
| ROT | 1 | 0,312 | 0,084 | 0,605 | BAL2 | 3.2 | 0,066 | 0,911 | 0,022 |
| CHS | 1.1 | 0,188 | 0,063 | 0,750 | BAL3 | 3.2 | 0,218 | 0,733 | 0,050 |
| WAS | 1.1 | 0,169 | 0,031 | 0,801 | BAL4 | 3.2 | 0,263 | 0,695 | 0,042 |
| AMS | 1.2 | 0,037 | 0,070 | 0,894 | RHE2 | 4 | 0,275 | **0,636** | 0,089 |
| KOS | 1.2 | 0,093 | 0,039 | 0,867 | MAI7 | 4.1 | 0,236 | 0,127 | **0,637** |
| STS | 1.2 | 0,036 | 0,026 | 0,937 | BOS1 | 4.2 | 0,037 | 0,059 | **0,904** |
| WLS | 1.2 | 0,021 | 0,015 | 0,963 | BOS3 | 4.2 | 0,027 | 0,019 | **0,954** |
| ELB6 | 2 | 0,752 | 0,043 | 0,205 | BOS4 | 4.2 | 0,166 | 0,043 | **0,791** |
| ELB7 | 2 | 0,714 | 0,155 | 0,131 | BOS5 | 4.2 | 0,150 | 0,044 | **0,806** |
| HAV1 | 2 | 0,793 | 0,097 | 0,110 | EDS | 5 | 0,345 | 0,204 | **0,451** |
| KAR | 2 | 0,801 | 0,104 | 0,095 | STM | 5.1 | 0,843 | 0,051 | 0,107 |
| JAG | 2 | 0,855 | 0,062 | 0,083 | EMS3 | 6 | 0,677 | 0,195 | 0,128 |
| KRK | 2 | 0,878 | 0,078 | 0,044 | EMS2 | 6 | 0,819 | 0,076 | 0,105 |
| STP | 2 | 0,713 | 0,142 | 0,144 | EID1 | 7 | 0,439 | 0,060 | **0,501** |
| MUR | 2.1 | 0,821 | 0,157 | 0,023 | UCK4 | 8 | 0,299 | 0,653 | 0,048 |
| SUS | 2.1 | 0,952 | 0,026 | 0,021 | HAH | 8.1 | 0,188 | 0,778 | 0,034 |
| GST | 2.1 | 0,837 | 0,054 | 0,108 | PEE | 9 | 0,183 | 0,732 | 0,085 |
| GKB | 2.1 | 0,527 | 0,375 | 0,097 | GPS | 10 | 0,270 | 0,059 | **0,671** |
| KDO | 2.1 | 0,649 | 0,308 | 0,043 | GRA | 11 | 0,439 | 0,494 | 0,068 |
| DRS | 2.1 | 0,802 | 0,078 | 0,120 | BAR | 12 | 0,169 | 0,743 | 0,088 |
| WIS | 2.2 | 0,317 | 0,095 | **0,588** |  |  |  |  |  |

\* See Table 1 for definition of sample IDs. Samples with the same number have been pooled for a clearer presentation in Fig. 4

Network analysis with mitochondrial *cyt b* sequences identified two of the three main haplotypes postulated by Skog et al. (2014). However, while mitochondrial haplotypes frequencies differed among drainage basins, there was not one to one correspondence of haplotypes with the main clusters identified here based on nuclear data. Of 184 sequences 159 (86.4%) grouped with five reference sequences defined as haplotype E, representing the northern clade of *E. lucius* (Fig. S3). 24 sequences (13.0%) grouped together with 16 haplotype B reference sequences, constituting the circumpolar clade, and one sequence grouped with three haplotype F reference sequences of the southern clade. The northern clade exhibited a star-like appearance consisting of the main haplotype, surrounded by 26 sub-haplotypes, deviating by one (N = 22) or two mutations (N = 4). Pike individuals from the North Sea hydrogeographic region were the dominant fraction (59.9%), while individuals from the Baltic and Black Sea regions contributed 30.2% and 10.7%, respectively. The circumpolar clade consisted mostly of pike from the Baltic Sea hydrogeographic region (87.5%) and only of a minority of pike from other regions (North Sea: 4.2%, Black Sea: 8.3%). It was separated from all but one (B14) haplotype B reference sequences by one mutation and appeared homogenous, except for two sub-haplotypes deviating by one mutation. The two clades (the main haplotypes E and B) differed by six mutations and were connected via a hypothetical ancestor. The closest representative of the southern clade, connected via the same ancestor, differed by four mutations from the northern and by six mutations from the circumpolar clade (Fig. S3).

### Population structure among the major hydrogeographic basins

Admixture analysis based on microsatellites, revealed varying proportions of the different genetic lineages in pike populations across hydrogeographic regions. The Black Sea genetic cluster was most frequent in pike of the Danube and its tributary rivers (65.3% in pie chart 1 of Fig. 4 – subsequently indicated as e.g. “65.3% in 1”) as well as in pike of the big alpine lakes (77.5% in 1.1 and 91.5% in 1.2) (Table 3). Elevated levels of the Black Sea lineage, however, were also found in the geographically close Lake Constance (86.4% in 4.2) and the river Main (63.7% in 4.1), a big tributary of the river Rhine connected via a channel – the Rhein-Main-Donau-Kanal – with the Danube. Interestingly, some water bodies in the very north of Germany hosted a high proportion of Black Sea genetic imprint as well, such as pike from the river Eider (50.1% in 7) and pike inhabiting Wittensee (58.8% in 2.2) and Großer Plöner See (67.1% in 10).

**Fig. 4:**
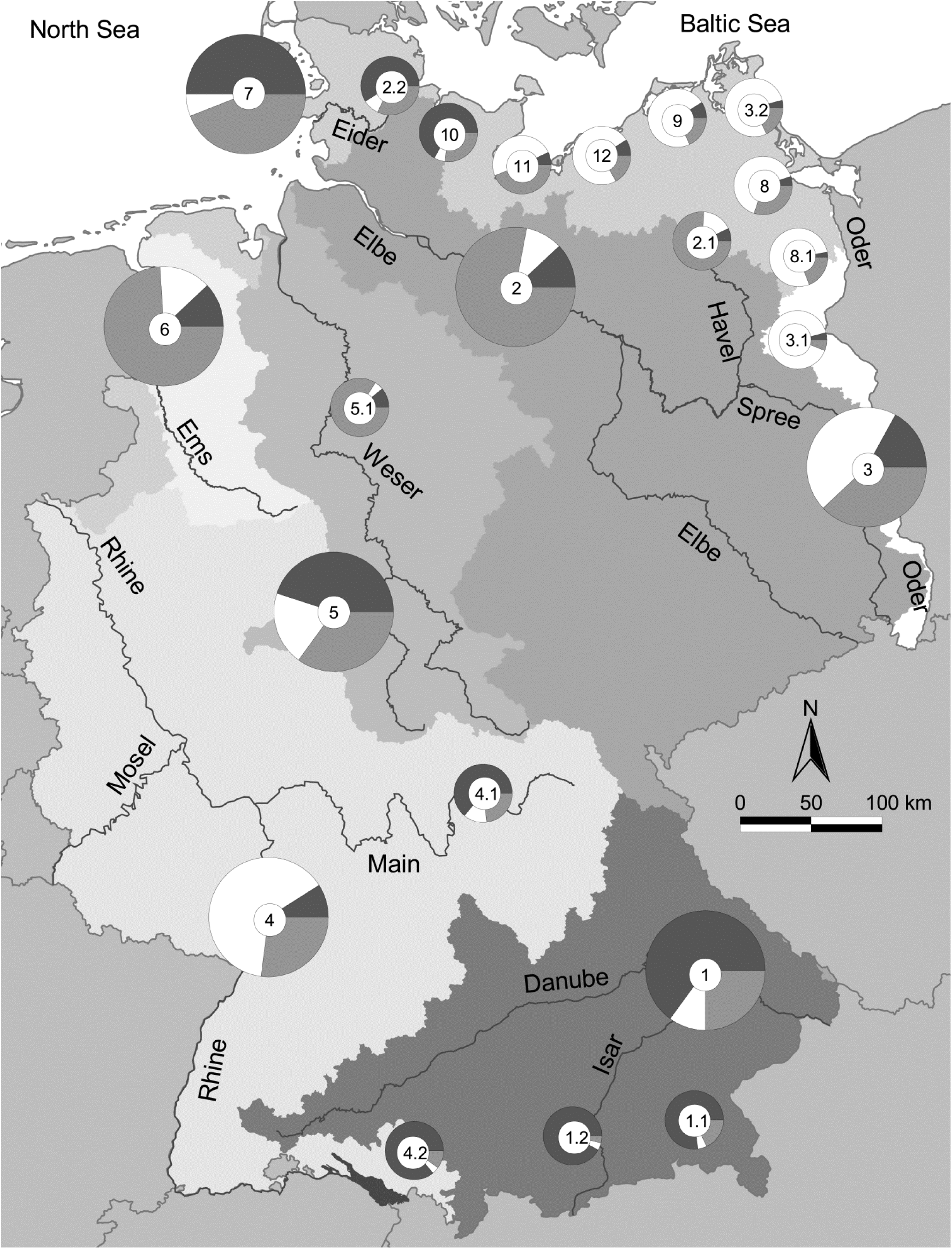
Map of Germany illustrating that genetic admixture on population level varied strongly and was not confined to a particular hydro-geographic region or river catchment therein. – Black, grey and white colors of the pie charts indicate genetic ancestry proportion of Black Sea, North Sea and Baltic Sea hydrogeographic region, respectively. Numbers indicate pooled populations as displayed in Table 3.

The Baltic genetic cluster was most prevalent in pike populations of the coastal waters of the Baltic Sea and water bodies draining into that basin, albeit with considerable variation (49.4% to 89.7% in 3.1, 3.2, 8, 8.1, 9, 11 and 12). Among the populations from Baltic tributaries, pike from the Oder main stream and from its tributary river Neiße, showed signs of pronounced admixture with the Black Sea and North Sea genetic lineages (in total 57.2% in 3).

The North Sea genetic cluster was dominant in populations of the river catchments of Elbe (78.7% in 2 and 76.5% in 2.1, excluding Wittensee with only 31.7% in 2.2), Ems (74.8% in 6) and Weser draining into the North Sea. Populations of the Weser catchment were represented by pike of two big lakes, Steinhuder Meer and Edersee (a reservoir), which differed in their admixture patterns. While populations of the Steinhuder Meer were predominantly shaped by the North Sea genetic cluster (84.3% in 5.1), this ancestry contributed relatively little to pike of the Edersee population (34.5% in 5). Similarly, and unexpectedly, the river Rhine exhibited a higher proportion of the Baltic than of the North Sea genetic cluster (63.6% in 4).

Genetic admixture could also be read from the distance based consensus tree (Fig. 2) and the frequency based principal coordinate analysis (Fig. 3). E. g., the Rhine population (RHE2) appeared within the Baltic Sea cluster and pike from Großer Plöner See (GPS) were placed within the Black Sea cluster.

### Admixture at the individual level

Genetic admixture was examined at the level of individuals within pike populations to assess the homogeneity of ancestries and investigate for possible signs of population substructure (Fig. 5). Based on ancestry coefficients (NA), the distributions of native *vs*. foreign genetic ancestries displayed a range from mostly pure native populations (green violin plots in Fig. 5 with mean NA ≥ 0.50) to hybrid swarms, with mostly admixed individuals (yellow violin plots in Fig. 5 with 0.25 < NA < 0.50). Moreover, some distributions were skewed towards foreign ancestry, with complete replacement of native ancestry in some populations (red violin plots in Fig. 5 with mean NA ≤ 0.25). The frequency distribution of ancestry coefficients in some populations showed signs of bimodality, that is, individuals may fall into different groups that differ in their ancestry coefficients (Fig. 5). This includes river populations of e.g. the Danube (DON, INN, NAB, ROT; see table 1 for explanation of IDs) and Oder catchments (ODE2, ODE7, NEI2) as well as lake populations of e.g. the Trave (GRA) and Elbe catchments (GKB). In other populations, such patterns were observed to a less extent, e.g. in lake populations of the Ucker (HAH) and Weser catchments (STM) and river populations of the Elbe catchment (HAV1, KAR).

**Fig. 5:**
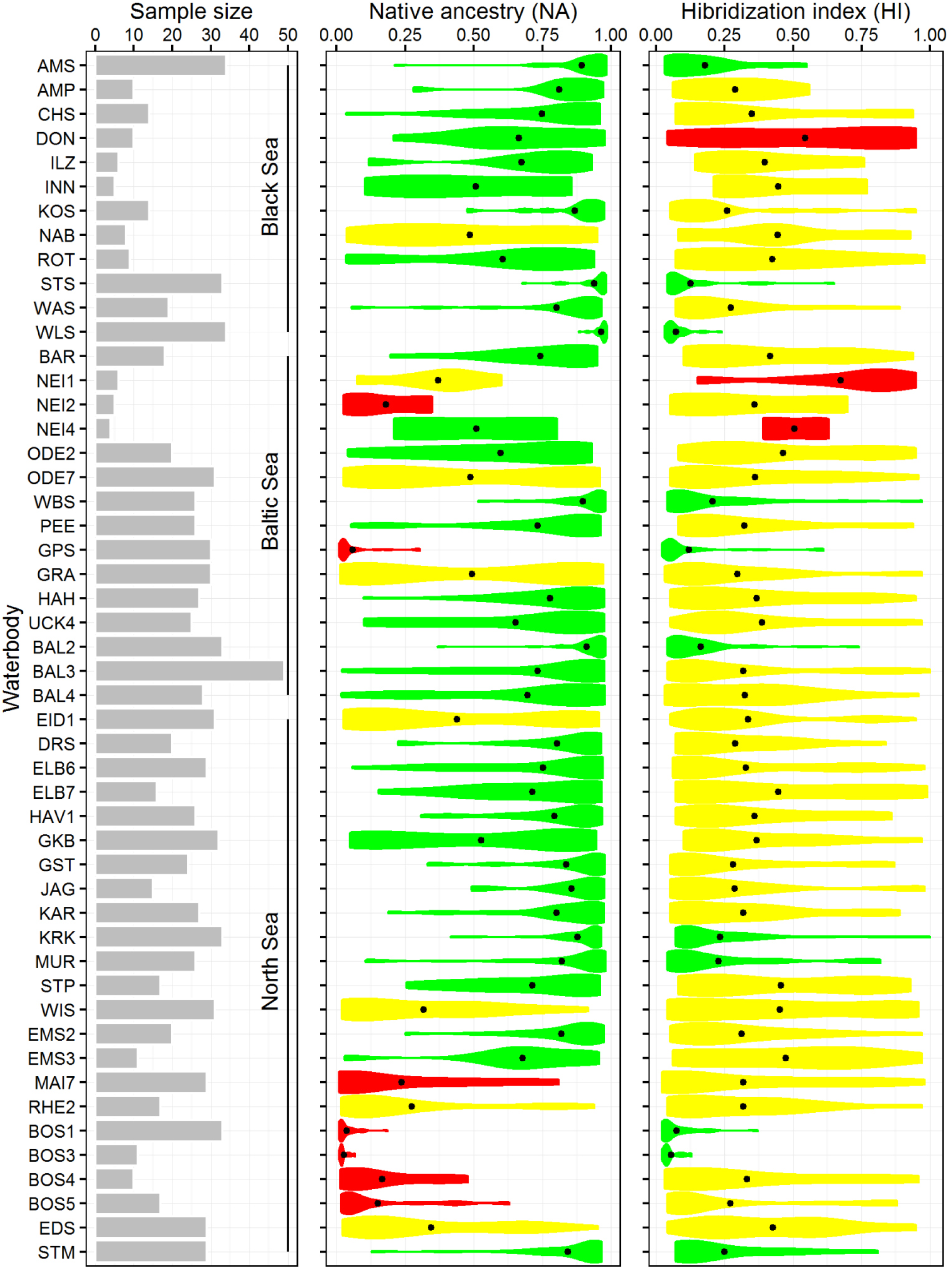
Genetic admixture calculated per individual. Extension of a figure indicates increased numbers of individuals with a certain proportin of native ancestry (column 2) or degree of hybridization (column 3). Mean values are indicated within each figure as a dot. Column 1 indicates the number of individuals analyzed per sampling site. Color code for native ancestry: green = NA ≥ 0.5, yellow = 0.25 < NA < 0.5, red = NA ≤ 0.25, color code for hybridization index: green = HI ≤ 0.25, yellow = 0.25 < HI < 0.5, red = HI ≥ 0.5.

Ancestry distributions within pike populations of the Black Sea hydrogeographic region generally exhibited higher proportions of native ancestries (particularly in pike of the alpine lakes), whereas the two other hydrogeographic regions were comprised of populations in which individual genotypes suggested high proportions of foreign genetic material. The most extreme examples included pike of Großer Plöner See (GPS) and of the rivers Neiße (NEI2) and Main (MAI), where a near complete replacement of native by foreign genetic identities was suggested by the most likely STRUCTURE model (Fig. 5) and confirmed by genetic distance trees (Fig. 2) and principal coordinate analysis (Fig. 3). Pike populations in Lake Constance, however, ought to be viewed differently in this regard due to their geological history (see discussion for further details). In the Baltic Sea hydrogeographic region, a coastal population (BAL2 in Fig. 5) and one freshwater population (WBS) exhibited pronounced native genetic signatures. Likewise, the North Sea hydrogeographic region harbored populations that appear to be rather typical and pure representatives of the respective genetic cluster (JAG, KRK, GST, MUR).

### Correlation of hybridization levels with ecological quality

Employing hierarchical general linear models revealed that the ecological status of the water body as well as the type of ecosystem had a significant effect on the hybridisation index (HI) of pike populations. Specifically, the decline in the ecological status was highly significantly correlated with HI (Table 4). Each unit of decrease of the ecological status lead to an increase of HI by a factor (slope) of 0.25 (1.28 in raw scale) ± 0.08 with respect to the intercept (defined as the best ecological status). Accordingly, the HI increased by a value of 0.21 from the best (= 1 in Fig. 6) to the poorest (= 5) ecological status defined according to the European Water Framework Directive. The estimate (in logit scale) of HI for lakes was −1.79 (0.17 in raw scale) ± 0.26, while it was −1.37 (0.26 in raw scale) ± 0.26 in rivers. These values indicated a significantly stronger signal of past hybridization of pike populations in rivers as compared to lakes (Table 4). While the hypothesis of a correlation between HI and the type and ecological status of the water body was supported, relationships of HI and the general level of modification of the water body (Table 4) and all two-level interaction effects were not supported (data not shown).

**Table 4:** Results of hierarchical general linear mixed modelling (HGLM) to test the effect of different linear predictors on the hybrid index (HI) and the native ancestry (NA) controlling for the random variance attributed to the individuals sampled in specific waterbodies nested within catchments (see Fig. 6). The table shows the estimates (in logit scale) and their standard error (s.e.), the t-value statistics and their p-value (Pr(>|t|). Two-level interactions were non-significant in all cases and removed from the model. The estimates of the categorical variables were shown per one category with respect to the other (intercept). Significance codes: 0 ‘***’ 0.001 ‘**’, 0.01 ‘*’, 0.05 and ‘.’ 0.1.

| Hybrid index | Estimate | s.e | t-value | $\Pr(>\vert t\vert )$ | |
| --- | --- | --- | --- | --- | --- |
| (Intercept) | -1.401 | 0.176 | -7.98 | <0.001 | *** |
| Type (river) | 0.376 | 0.119 | 3.16 | <0.01 | * |
| Modification (high) | -0.128 | 0.125 | -1.03 | 0.3 |  |
| Ecological Status | 0.186 | 0.053 | 3.49 | <0.001 | *** |

| Native ancestry | Estimate | s.e | t-value | $\Pr(>\vert t\vert )$ | |
| --- | --- | --- | --- | --- | --- |
| (Intercept) | 0.731 | 0.44 | 1.66 | 0.096 |  |
| Type (river) | -0.247 | 0.239 | -1.030 | 0.301 |  |
| Modification (high) | 0.389 | 0.266 | 1.460 | 0.143 |  |
| Ecological Status | -0.16 | 0.104 | -1.550 | 0.122 |  |

**Fig. 6:**
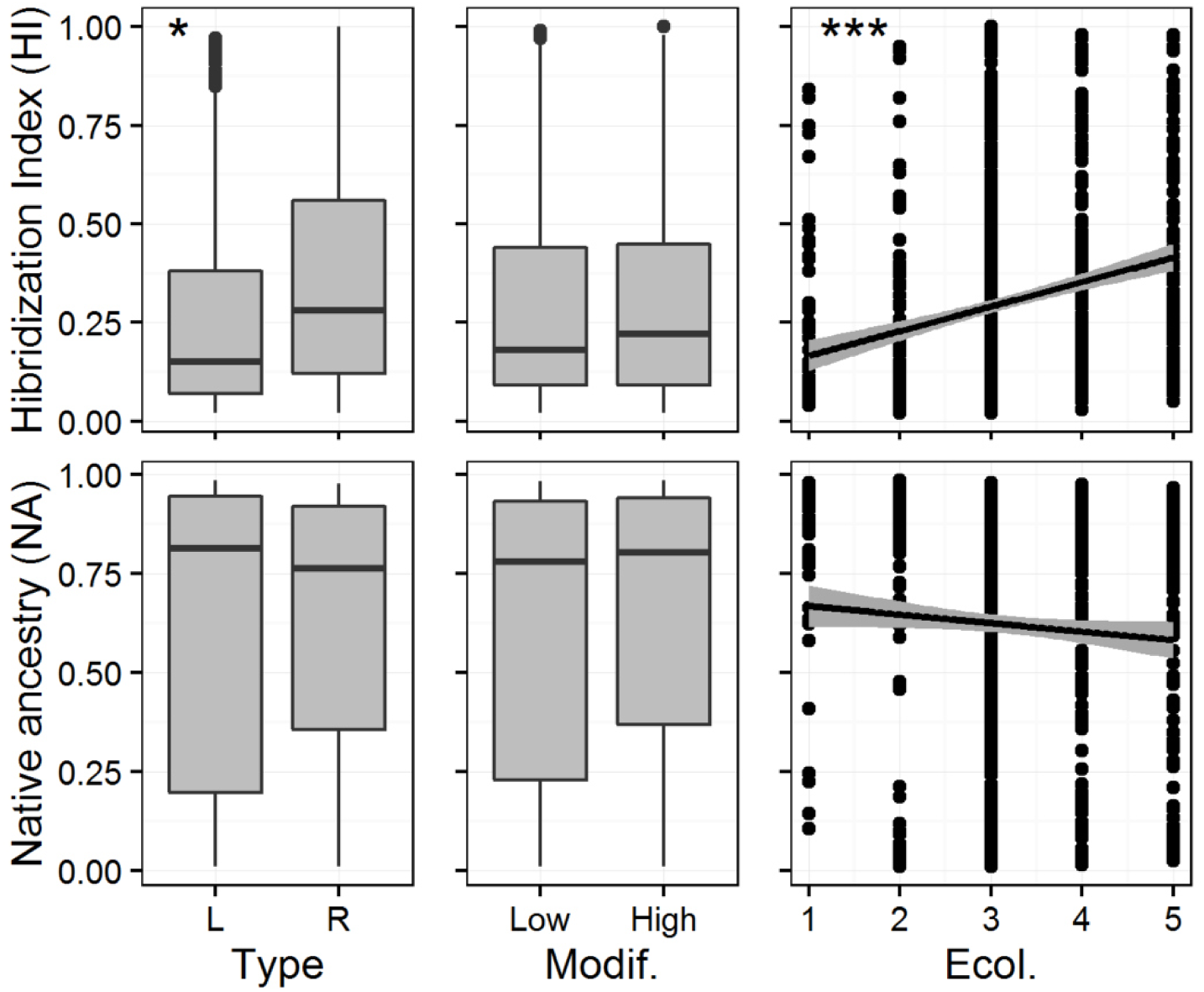
Correlation of native ancestries and hybridization indices with habitat type, strength of modification and ecological quality (1 = very good to 5 = poor according to EU water framework directive) as obtained with HGLM analysis (see Table 4 for details). Baltic coastal waters (BAL2, BAL3 and BAL4) and freshwaters without ecological information (AWU, SUS, KDO) were excluded from analysis. IDs of water bodies are explained in Table 1.

In addition, the native ancestry (NA) exhibited a strong negative correlation with the deterioration of the ecological status of the water bodies (Fig. 6), however this was not statistically significant (Table 4). This held also true for the predictors “type” (i.e. ecosystem type) and “modification” (i.e. general degree of modification) of the water bodies (Table 4), as well as their two-level interactions (not shown), which were therefore removed from the model with NA.

## Discussion

### Differentiation of pike from different drainages

Mitochondrial DNA markers have been previously used to reconstruct the colonization patterns of pike in Europe after the last glaciations c. 15.000 years ago (Nicod et al., 2004; Skog et al., 2014). Our own analysis of mitochondrial *cyt b* sequences together with 15 polymorphic microsatellites (Eschbach & Schöning, 2013) allowed distinguishing lineages that seem typical for different drainage basins. STRUCTURE analysis (Falush et al., 2003) of microsatellite data revealed a k value of three as the most likely number of genetic lineages present in Germany (Fig. 1). A second k value of five was deemed much less likely when judged against results obtained with two other analysis performed with microsatellite data. Both the construction of genetic distance based consensus trees as well as a principal coordinate analysis based on allele frequencies argued in favor of only three genetic clusters. Each of them was regarded as representative of the hydrogeographic regions in the North Sea, Baltic Sea or Black Sea, suggesting that these were the most likely distribution areas of the ancestors after the retreat of the glaciers (Table 3). NETWORK analysis (Bandelt et al., 1999) of *cyt b* haplotypes assigned about 90% of pike originating from the hydrogeographic region of the Baltic Sea to the circumpolar clade and almost 60% of pike originating from the North Sea region to the northern clade as described by Skog et al. (2014). In our data the southern clade, identified as a third mitochondrial haplotype by Skog et al. (2014), was represented by only a single individual among 21 pike from the Danube catchment (Fig. S3). Thus, although our analysis of mitochondrial haplotypes agrees with the general findings from Skog et al. (2014), our data showed that lineage sorting of mitochondrial haplotypes has not proceeded to a point where haplotypes alone are sufficient to distinguish the lineages of pike studied here. Hence, the strongest support for the existence of three evolutionarily significant units (Moritz, 1994) of pike with different distribution areas was supported by multilocus microsatellite analyses.

### Signatures of migration or stocking?

Recent secondary contacts and genetic admixture between divergent pike lineages have most likely increased as a result of anthropogenic activities. This included natural migration through human-made artificial connections among different river basins as well as stocking of economically important fish species. The latter represents an important factor that increases the potential for gene flow between populations naturally separated in space. Unfortunately, past stocking is generally not well documented in Germany (Arlinghaus et al., 2015), but certainly has occurred over decades in central Europe in pike and multiple other economically relevant fishes (Cowx, 1994; Guillerault et al., 2018; Kottelat & Freyhof, 2007; Larsen & Berg, 2004). Given the poor locality-specific records, it is however impossible to take stocking into account in a more detailed way than just to accept that it has happened frequently across pike stocks (Arlinghaus et al., 2015). Nevertheless, the pike system offers the opportunity to explore environmental factors that stabilize existing diversity patterns or, conversely, promote admixture, without knowing which factor of secondary contact was ultimately responsible. Pike of the Danube catchment and of the brackish coastal areas of the Baltic Sea exhibited a clear dominance of native ancestry as inferred from analysis of 15 polymorphic microsatellites (Fig. 4). Likewise, relatively low admixture levels were observed in pike populations of the Ems (pie-chart no. 6 in Fig. 4) and Weser catchments (pie-chart no. 5.1 in Fig. 4) belonging to the North Sea hydrogeographic region. A possible explanation for the persistence of autochthonous populations is either a low level of local stocking or competitive exclusion of foreign genotypes by better-adapted native populations (Englbrecht et al., 2002; Eschbach et al., 2014; Gandolfi et al., 2017). It has been found before that brackish water populations are adapted to reproduction in low salinity conditions, causing an increase in the mortality of stocked freshwater fish, thereby preventing introgression despite decades of stocking (Jørgensen et al., 2010; Larsen et al., 2005). Moreover, high density blocking can effectively counteract establishment of immigrants from a distant population in an environment already inhabited by locally adapted conspecifics as long as the local stock is naturally reproducing at high levels (van Poorten et al., 2011; Waters, 2011). In line with this, stocking experiments with pike showed that stocked individuals suffer from substantially higher mortality than wild conspecifics, if the natural reproduction is sufficient, reducing the potential for successful establishment (Arlinghaus et al., 2015; Diana et al., 2017; Hühn et al., 2014).

In some water bodies investigated in this study, however, pike individuals showed high foreign ancestries, e.g. in the rivers Neiße, Main, Rhine and Eider, as well as in the lakes Wittensee and Edersee. This data suggested that much of the genetic material from native populations may be replaced through the introduction of foreign stocks, which in turn can maintain populations that are resilient to further invasion by local genotypes. In this context, our analysis suggests a near complete replacement of native pike in the lake Großer Plöner See, in the north of Germany that is presently inhabited by pike characterized by microsatellite genotypes that are likely to originate from the Danube catchment. Admittedly, the data set we used in this study is heterogeneous and complex, which makes an analysis of the overall population structure a difficult task. Still, sampling for this particular lake is above average (N = 30 individuals), and the effect we observed according to the most likely STRUCTURE model was unambiguous and supported by two other analytical methods (genetic distance based consensus trees and allele frequency based PCoA). Englbrecht et al. (2002) described a comparable case for the arctic char (*Salvelinus umbla*) in Starnberger See (Bavaria, Germany), where resident fish have been completely replaced by stocked fish. They argue that this was possible because the lake became heavily eutrophied in the middle of the last century (Ruecker et al., 1999), with resident char approaching near extinction, while non-resident char were apparently adapted to deal with the novel environment. There is clear evidence that such a complete replacement has occurred in a range of other stock-enhanced populations of fishes (van Poorten et al., 2011). Thus, although no detailed ecological data was available for Großer Plöner See, it is possible that its native population might have undergone a similar fate and became invaded by stocked pike. This example shows that complete genetic swamping most likely by stocking is indeed a possible scenario even if the ecosystem appears healthy and of high integrity in present time. By contrast, the high proportion of Black Sea hydrogeographic region ancestry in pike of Lake Constance is likely a result of ancient natural connections with the Danube catchment and rather reflects native ancestry stemming from natural post-glacial dispersal, previously reported for perch (*Perca fluviatilis*) by Behrmann-Godel et al. (2004) for the same system. The minor proportions of other non-North Sea hydrogeographic region ancestries were likely due to human-assisted colonization via introduction and stocking, because southward gene flow from downstream areas of the river Rhine is not possible due to an insurmountable waterfall at Schaffhausen (Switzerland).

Pike populations in other water bodies exhibited high levels of genetic admixture. When source populations are adjacent, admixture can be explained by natural immigration through man-made connections such as the “Main-Donau Kanal” linking the Danube with the river Main (Powels et al., 2013). However, we also detected signatures of admixture between rather distant source populations, e.g. between pike of the rivers Oder in the east and Rhine in the west or between the rivers Danube in the south and Eider in the very north of Germany. Stocking, rather than migration, is a more likely explanation here, because migration would probably have created a more coherent geographical pattern. Our data are in line with genetic structures of pike populations in Denmark at the intra-specific level (Bekkevold et al., 2015) and Italy at the inter-specific level (Gandolfi et al., 2017), both of which not always reflected natural catchment barriers and were likely caused by successful pike stock enhancement activities in the past.

Pike of the lakes Großer Kossenblatter See, Drewitzer See and Ammersee exhibited pronounced linkage disequilibria (Fig. S2), and all pike of lake Ammersee additionally exhibited deviation from Hardy-Weinberg equilibrium as well as heterozygote deficiencies (Fig. S1). In agreement with this, bimodal distributions of native and foreign ancestries of individuals of some populations confirmed that they were not genetically homogenous. This can be compared to bimodal hybrid zones, which are often characterized by pronounced deviation from Hardy-Weinberg equilibrium due to restrictions in panmixia (Allendorf et al., 2001; Redenbach & Taylor, 2003). Possible scenarios to explain this pattern include that foreign pike genotypes are regularly introduced at a large scale without much reproductive success. Alternatively, foreign genotypes may be reproductively isolated to some extent so that they persist as a distinct genetic population in parallel to the local population of pike. The latter explanation is less likely because we know that stocked individuals that survive readily hybridize with native pike (Arlinghaus et al., 2015), although there is evidence of natal homing of pike in large standing water bodies, which can contribute to the development of meta-populations within lake ecosystems (Miller et al., 2001).

### Impacts of ecosystem status on hybridization

In other cases, we detected various degrees of foreign ancestry (Fig. 5), documenting likely admixture between genetically distinct populations – an effect that increased with the degradation of the ecological status of the recipient ecosystem. We note that the quality of this inference depends on the sample sizes that were available for each population as well as the degree of differentiation between the presumed source populations. It might, therefore, be useful to revisit specific populations with a more powerful study design and genome wide marker coverage. Nonetheless, it was obvious that the three pike lineages readily hybridized upon secondary contact. This result bears general questions on why hybridization proceeded with different intensity in different pike populations and whether pike of different origins are indeed isolated to some extent when they are brought into secondary contact.

We found that the individual admixture levels in pike, expressed as a hybridization index (HI), were not confined to a specific hydrogeographic region or any particular river catchment therein. Instead it turned out that the HI increased significantly with decreasing ecological quality of a water body. Albeit not statistically significant, we observed a congruent decrease of native ancestry. Thus, environmental change could have driven genetic changes in pike populations and individuals by affecting the frequency of hybridization among populations brought into secondary contact. The fact that the HI was only slightly lower in water bodies with low modifications as compared to the HI of highly modified waters demonstrates that the admixture as such occurs in all populations and is not restricted to highly modified habitats (Fig. 6).

Our analysis yielded a significantly higher HI in pike populations in rivers as compared to lake-dwelling pike, which is likely due to fundamental ecological differences between the two habitat types such as the increased natural connectivity in rivers, resource availability, productivity, habitat structure, and community composition (Irz et al., 2006; Hof et al., 2008). Most importantly, however, rivers and lakes vary in stability and disturbance frequency, including exposure to catastrophic floods, which occur more frequently in lotic than in lentic systems. Rivers of central Europe also have been more strongly modified, e.g. by removal of connectivity to floodplains and habitat simplification, which represent a central component of their disturbance regime and at the same time constitute essential spawning habitat for pike. In a recent meta-analysis comparing resistance of limnic, marine and terrestrial ecosystems towards invasive species, Alofs & Jackson (2014) clearly demonstrated that lentic systems displayed a higher biotic resistance than lotic systems, which is in accordance with our findings of different susceptibilities towards hybridization in river and lake pike populations.

Our observation that hybridization in pike appears to be favored in ecologically perturbed water bodies raises important questions about the mechanisms. The effect could first be caused due to an increase of foreign genotypes that managed to invade a weakened native population (Englbrecht et al., 2002; Gandolfi et al., 2017). Alternatively, genetically admixed fish could be more competitive in the face of anthropogenic changes to the ecosystem. This would resemble the first step of a hybrid speciation scenario, where intraspecific hybrids are expected to be most successful when parental populations are not at their optimum (Abbott *et al.*, 2013; Nolte & Tautz, 2010). Stelkens *et al*. (2014) showed that particularly the interactions of genetic variants between distant *Saccharomyces* strains can lead to a better survival in environments of decreasing quality. Thus, hybridization can create biodiversity resulting in novel phenotypes and adaptive change in response to environmental change (Arnold, 2016; Charlesworth & Willis, 2009; Edmands, 2007; Sefc et al., 2017). Examples of these processes can be found among invaders conquering new environments that were not occupied by populations of the respective species before, as it was found for *Cottus* hybrids in the river Rhine (Nolte et al., 2005; Stemshorn et al., 2011), but also for spiders (Krehenwinkel & Tautz, 2013) and some plants (Keller & Taylor, 2010). Likewise, in a previous study we observed increased intraspecific genetic diversity of zander (*Sander lucioperca*) in water bodies, where this fish species had been introduced in the late 19^th^ century, a pattern that would be in line with an advantage of admixed individuals in the course of an invasion (Eschbach et al., 2014). Thus, new combinations of genes from different evolutionary backgrounds might enable fast adaptation, and thus increase the chance to survive under worsening environmental conditions (Arnold 2016). However, careful future studies are needed to distinguish the adaptive scenario outlined here from neutral explanations that are related to abrupt changes in propagule pressure in fluctuating environments.

### Conclusions and implications

At the species level, large-scale hybridization, which extends over different hydrogeographic regions each with its own evolutionary history of genetic lines, is synonymous with genetic erosion (Epifanio & Philipp, 2001), i.e. it increases the fate of extinction due to the loss of evolutionary potential. Especially in a species such as pike, which is characterized by a low natural genetic variability compared to other freshwater fish, it may be important to maintain genetic diversity through different genetic lines. Our study clearly showed a novel relationship between ecosystem status, assessed under the European Water Framework Directive, and the genetic structure of northern pike. It supports the idea that habitat degradation can also have far-reaching consequences for genetic integrity within species and promotes efforts to further improve the ecological quality of lakes and rivers. In the case of pike, this would essentially mean reconnecting floodplains with rivers and reducing nutrient inputs into lakes, which would increase both population size and genetic biodiversity.

## Acknowledgements

Particular thanks go to Yvonne Klaar, Asja Vogt, Jasmin Spamer and Elke Bustorf for technical assistance in the lab, as well as Petra Kersten for invaluable advice on microsatellite analysis. We are also indebted to Ute Mischke for kindly providing additional ecological data of the different water bodies. We thank Jochem Kail for creating the map in Fig. 4. We are very much obliged to the many fishers, anglers, fisheries authorities and research organisations for providing tissue samples as well as a lot of helpful information. We are grateful to the whole BESATZFISCH team and Michael Monaghan and his POPGEN group for many fruitful discussions. Funding of the current work was granted by the German Federal Ministry of Education and Research within the project BESATZFISCH (www.besatz-fisch.de) in the Program for Social-Ecological Research (Grant No. 01UU0907 to R. A.). A.N. was supported through an ERC starting grant “EVOLMAPPING”.

## Legends to supplementary material

**Fig. S1:** Pike populations showing deviations from Hardy-Weiberg equilibrium (A) and heterozygote deficiency (B).

**Fig. S2:** Number of loci combinations exhibiting linkage disequilibria determined with and without Bonferroni correction.

**Fig. S3:** Network analysis based on *cyt* b sequences. B, E and F mark the circumpolar, northern and southern clades, respectively, of the northern pike according to Skog *et al*. (2014). – The size of the circles is proportional to the number of pike individuals with a certain haplotype; color code for samples: blue = North Sea, green = Baltic Sea, orange = Black Sea hydro-geographic region, gray = reference sequences, white = a mutation step, black = a hypothetical ancestor.

**Table S1:** Test for evidence of null alleles. **A**: number of alleles per locus and population (FSTAT 2.9.3.2). Total number of alleles = 6,969. **B**: Test for null alleles (MICROCHECKER 2.2.3). Total number of putative null alleles = 112, i.e. 1.6% of total number of alleles.

**Table S2:** Pairwise Fst (below diagonal) and p values (above diagonal) of 53 pike populations. Fst values were employed for principal coordinate analysis (Fig. 3).

