## supplemental figure 1 for "Intraspecific population admixture of a top piscivore correlates with anthropogenic alteration of freshwater ecosystems"

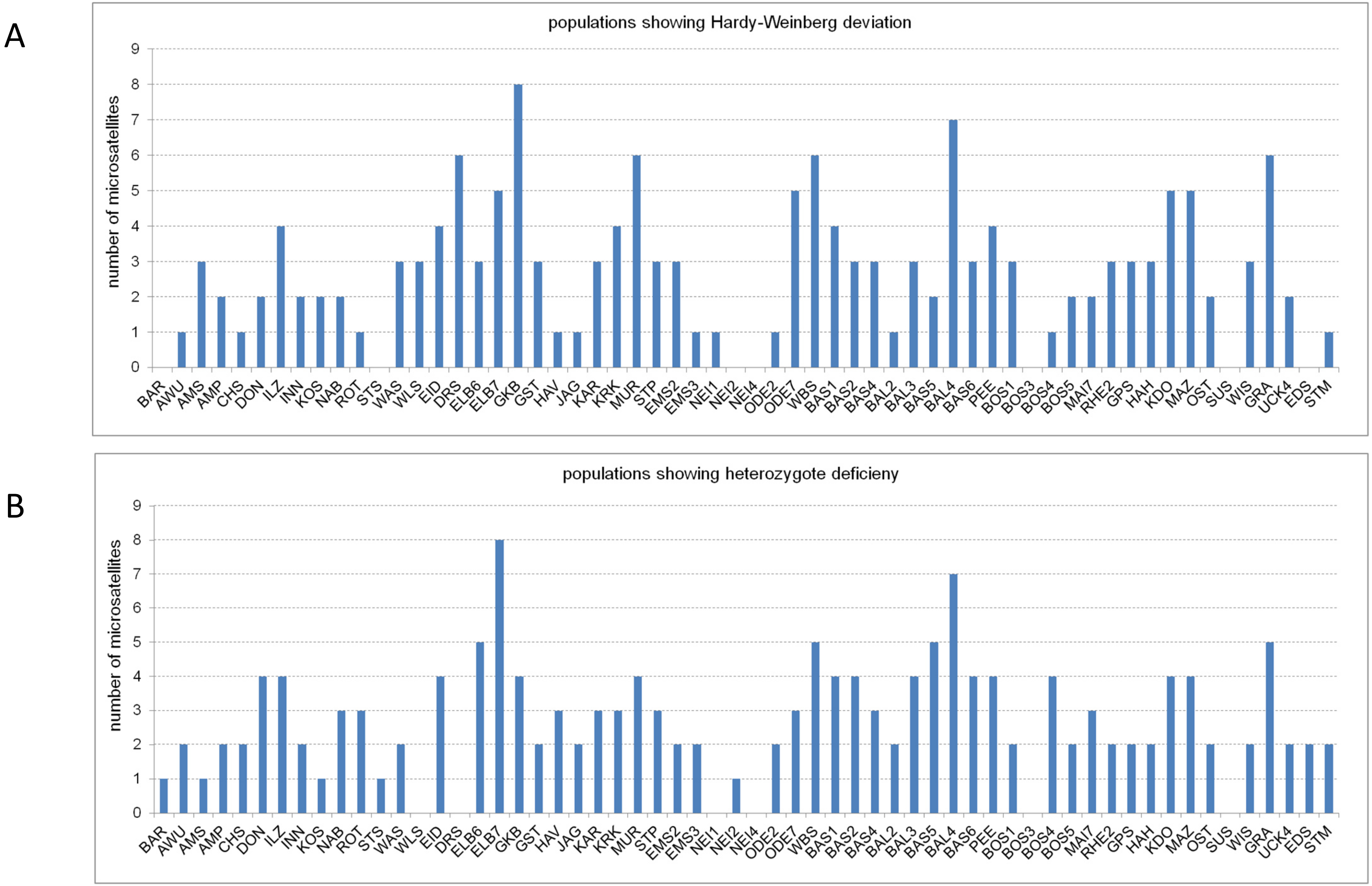


**Fig. S1**: Pike populations showing deviations from Hardy-Weiberg equilibrium (A) and heterozygote deficiency (B).
