## supplemental figure 2 for "Intraspecific population admixture of a top piscivore correlates with anthropogenic alteration of freshwater ecosystems"

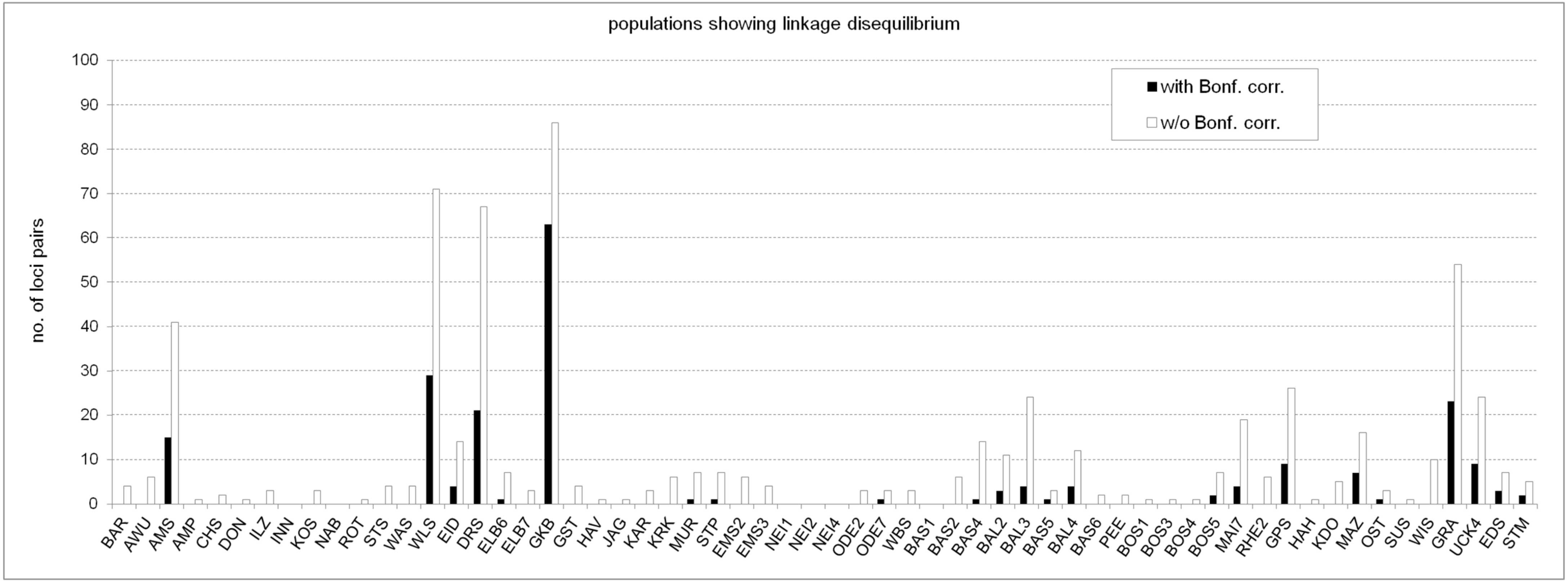


**Fig. S2**: Number of loci combinations exhibiting linkage disequilibria determined with and without Bonferroni correction.
