## supplemental figure 3 for "Intraspecific population admixture of a top piscivore correlates with anthropogenic alteration of freshwater ecosystems"

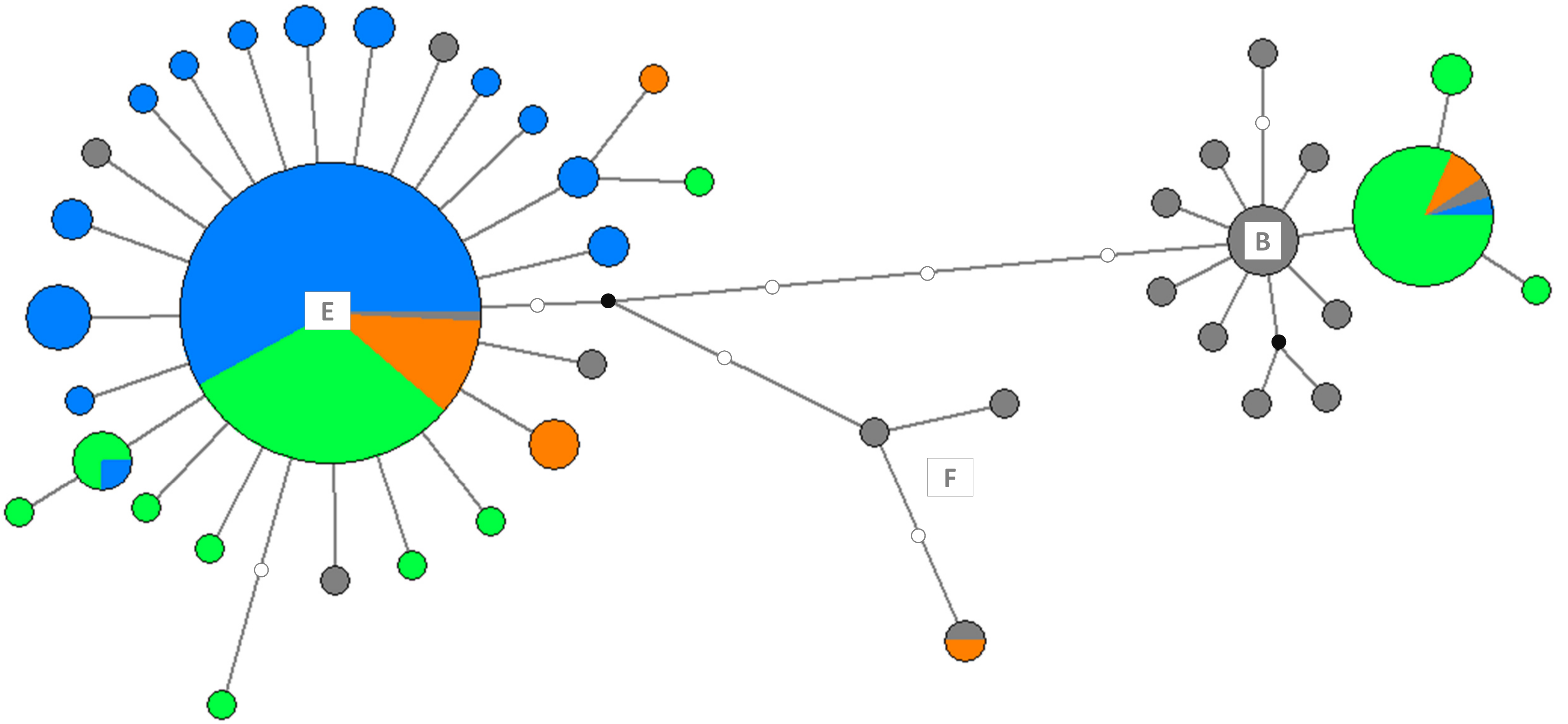


**Fig. S3**: Network analysis based on *cyt* b sequences. B, E and F mark the circumpolar, northern and southern clades, respectively, of the northern pike according to Skog et al. (2014). – The size of the circles is proportional to the number of pike individuals with a certain haplotype; color code for samples: blue = North Sea, green = Baltic Sea, orange = Black Sea hydro-geographic region, gray = reference sequences, white = a mutation step, black = a hypothetical ancestor.
