## supplemental table 1 for "Intraspecific population admixture of a top piscivore correlates with anthropogenic alteration of freshwater ecosystems"

**Table S1:** Test for evidence of null alleles. **A:** number of alleles per locus and population (FSTAT 2.9.3.2). Total number of alleles = 6969. **B**: Test for null alleles (MICROCHECKER 2.2.3). Total number of putative null alleles = 112, i.e. 1.6% of total number of alleles.

## A

| loci | BAR | AWU | AMS | AMP | CHS | DON | ILZ | INN | KOS | NAB | ROT | STS | WAS | WLS | EID |
|---|---|---|---|---|---|---|---|---|---|---|---|---|---|---|---|
| Elu87 | 3 | 4 | 9 | 5 | 6 | 7 | 4 | 3 | 5 | 3 | 4 | 7 | 7 | 4 | 3 |
| Eluc045 | 6 | 4 | 10 | 8 | 8 | 6 | 5 | 6 | 10 | 8 | 6 | 8 | 8 | 8 | 8 |
| B451 | 15 | 11 | 19 | 11 | 11 | 11 | 7 | 6 | 15 | 11 | 11 | 13 | 11 | 11 | 15 |
| PkB47 | 5 | 5 | 7 | 7 | 8 | 5 | 5 | 4 | 5 | 6 | 7 | 8 | 6 | 8 | 5 |
| Elu19 | 6 | 4 | 6 | 3 | 4 | 5 | 4 | 6 | 6 | 4 | 4 | 8 | 4 | 4 | 4 |
| El02 | 7 | 5 | 11 | 6 | 8 | 7 | 6 | 2 | 6 | 4 | 6 | 8 | 5 | 5 | 7 |
| Pkb16 | 15 | 6 | 15 | 11 | 11 | 10 | 7 | 7 | 12 | 11 | 10 | 16 | 16 | 13 | 18 |
| Elu76 | 6 | 7 | 8 | 7 | 9 | 8 | 5 | 4 | 7 | 5 | 7 | 8 | 7 | 5 | 10 |
| EL27 | 4 | 6 | 6 | 4 | 5 | 6 | 5 | 3 | 5 | 6 | 6 | 5 | 6 | 4 | 8 |
| EmaD12a | 14 | 8 | 13 | 10 | 11 | 13 | 7 | 7 | 14 | 9 | 7 | 14 | 12 | 10 | 13 |
| EL01 | 4 | 6 | 10 | 7 | 8 | 7 | 5 | 3 | 4 | 6 | 7 | 7 | 6 | 6 | 8 |
| EluB108 | 5 | 2 | 4 | 4 | 6 | 5 | 4 | 3 | 5 | 5 | 3 | 4 | 3 | 4 | 5 |
| EluBe | 3 | 2 | 5 | 5 | 4 | 4 | 3 | 3 | 3 | 2 | 3 | 3 | 3 | 4 | 3 |
| B24 | 8 | 8 | 12 | 9 | 10 | 11 | 7 | 6 | 13 | 11 | 10 | 16 | 11 | 11 | 12 |
| Eluc033 | 10 | 4 | 12 | 7 | 6 | 10 | 3 | 2 | 6 | 4 | 6 | 7 | 9 | 7 | 7 |

| loci | DRS | ELB6 | ELB7 | GKB | GST | HAV | JAG | KAR | KRK | MUR | STP | EMS2 | EMS3 | NEI1 | NEI2 |
|---|---|---|---|---|---|---|---|---|---|---|---|---|---|---|---|
| Elu87 | 5 | 7 | 5 | 6 | 6 | 5 | 5 | 7 | 6 | 5 | 4 | 4 | 3 | 3 | 3 |
| Eluc045 | 4 | 6 | 4 | 7 | 5 | 6 | 4 | 5 | 4 | 4 | 7 | 6 | 4 | 3 | 2 |
| B451 | 6 | 13 | 13 | 15 | 15 | 21 | 16 | 19 | 20 | 8 | 12 | 7 | 9 | 8 | 9 |
| PkB47 | 4 | 5 | 9 | 10 | 9 | 9 | 7 | 11 | 7 | 7 | 6 | 10 | 5 | 5 | 5 |
| Elu19 | 3 | 5 | 4 | 3 | 6 | 4 | 4 | 7 | 5 | 4 | 4 | 4 | 2 | 2 | 2 |
| El02 | 5 | 9 | 10 | 12 | 9 | 1 | 10 | 11 | 8 | 13 | 6 | 6 | 7 | 6 | 4 |
| Pkb16 | 13 | 17 | 17 | 22 | 16 | 20 | 14 | 21 | 21 | 18 | 11 | 16 | 16 | 8 | 6 |
| Elu76 | 5 | 5 | 8 | 7 | 7 | 7 | 6 | 9 | 8 | 6 | 6 | 8 | 6 | 5 | 6 |
| EL27 | 6 | 9 | 5 | 5 | 5 | 9 | 7 | 8 | 9 | 6 | 8 | 7 | 4 | 4 | 5 |
| EmaD12a | 11 | 15 | 15 | 14 | 13 | 14 | 10 | 16 | 13 | 14 | 10 | 13 | 8 | 9 | 8 |
| EL01 | 5 | 7 | 7 | 8 | 6 | 5 | 7 | 6 | 7 | 4 | 9 | 5 | 5 | 4 | 3 |
| EluB108 | 6 | 7 | 7 | 6 | 5 | 7 | 3 | 7 | 6 | 6 | 4 | 6 | 5 | 5 | 6 |
| EluBe | 2 | 4 | 4 | 4 | 3 | 5 | 4 | 4 | 3 | 3 | 4 | 3 | 3 | 3 | 3 |
| B24 | 9 | 12 | 8 | 12 | 13 | 10 | 11 | 11 | 14 | 10 | 11 | 12 | 8 | 7 | 4 |
| Eluc033 | 5 | 7 | 7 | 10 | 10 | 10 | 7 | 8 | 8 | 8 | 3 | 6 | 6 | 5 | 4 |

| loci | NEI4 | ODE2 | ODE7 | WBS | BAS1 | BAS2 | BAS4 | BAL2 | BAL3 | BAS5 | BAL4 | BAS6 | PEE | BOS1 | BOS3 |
|---|---|---|---|---|---|---|---|---|---|---|---|---|---|---|---|
| Elu87 | 5 | 5 | 5 | 5 | 4 | 5 | 5 | 5 | 5 | 5 | 5 | 6 | 6 | 6 | 5 |
| Eluc045 | 4 | 10 | 10 | 9 | 8 | 12 | 5 | 6 | 4 | 6 | 7 | 10 | 10 | 9 | 6 |
| B451 | 6 | 20 | 18 | 14 | 11 | 15 | 15 | 21 | 16 | 16 | 20 | 14 | 19 | 17 | 12 |
| PkB47 | 2 | 7 | 7 | 5 | 7 | 7 | 5 | 6 | 6 | 3 | 5 | 7 | 8 | 8 | 5 |
| Elu19 | 2 | 5 | 6 | 5 | 4 | 7 | 6 | 7 | 4 | 7 | 7 | 6 | 4 | 5 | 4 |
| El02 | 4 | 1 | 6 | 8 | 8 | 13 | 8 | 3 | 14 | 12 | 13 | 7 | 11 | 6 | 4 |
| Pkb16 | 7 | 18 | 21 | 15 | 10 | 17 | 14 | 16 | 16 | 13 | 21 | 13 | 17 | 20 | 13 |
| Elu76 | 5 | 8 | 13 | 7 | 8 | 11 | 5 | 9 | 9 | 5 | 10 | 9 | 9 | 6 | 3 |
| EL27 | 3 | 8 | 7 | 5 | 6 | 5 | 5 | 7 | 6 | 6 | 6 | 4 | 6 | 5 | 3 |
| EmaD12a | 5 | 16 | 18 | 16 | 12 | 22 | 10 | 21 | 14 | 17 | 15 | 12 | 16 | 17 | 11 |
| EL01 | 2 | 9 | 9 | 7 | 8 | 11 | 4 | 12 | 7 | 10 | 11 | 5 | 8 | 6 | 4 |
| EluB108 | 5 | 7 | 10 | 5 | 6 | 7 | 6 | 5 | 5 | 4 | 6 | 6 | 6 | 4 | 4 |
| EluBe | 3 | 3 | 5 | 3 | 3 | 4 | 3 | 3 | 3 | 4 | 4 | 3 | 3 | 5 | 4 |
| B24 | 6 | 12 | 13 | 10 | 6 | 14 | 8 | 12 | 11 | 10 | 11 | 11 | 11 | 19 | 11 |
| Eluc033 | 5 | 7 | 10 | 9 | 8 | 9 | 3 | 9 | 3 | 7 | 7 | 7 | 9 | 8 | 4 |

| loci | BOS4 | BOS5 | MAI7 | RHE2 | GPS | HAH | KDO | MAZ | OST | SUS | WIS | GRA | UCK4 | EDS | STM |
|---|---|---|---|---|---|---|---|---|---|---|---|---|---|---|---|
| Elu87 | 4 | 6 | 8 | 3 | 3 | 6 | 3 | 4 | 5 | 4 | 6 | 4 | 5 | 5 | 5 |
| Eluc045 | 5 | 7 | 11 | 3 | 8 | 7 | 4 | 14 | 11 | 3 | 6 | 5 | 8 | 8 | 6 |
| B451 | 4 | 9 | 18 | 4 | 19 | 19 | 11 | 25 | 8 | 8 | 15 | 15 | 14 | 8 | 11 |
| PkB47 | 2 | 5 | 7 | 3 | 4 | 6 | 2 | 11 | 4 | 2 | 6 | 2 | 6 | 10 | 10 |
| Elu19 | 3 | 3 | 6 | 3 | 3 | 7 | 3 | 5 | 5 | 1 | 4 | 4 | 4 | 4 | 3 |
| El02 | 3 | 6 | 13 | 6 | 8 | 7 | 11 | 14 | 6 | 7 | 7 | 13 | 12 | 6 | 10 |
| Pkb16 | 11 | 14 | 21 | 11 | 16 | 15 | 12 | 16 | 15 | 8 | 15 | 17 | 12 | 19 | 20 |
| Elu76 | 2 | 5 | 9 | 6 | 7 | 9 | 8 | 12 | 10 | 3 | 8 | 8 | 7 | 11 | 9 |
| EL27 | 3 | 5 | 7 | 4 | 7 | 9 | 6 | 7 | 6 | 5 | 4 | 7 | 7 | 6 | 7 |
| EmaD12a | 11 | 10 | 15 | 7 | 16 | 19 | 16 | 17 | 16 | 7 | 14 | 12 | 12 | 16 | 16 |
| EL01 | 4 | 6 | 7 | 3 | 8 | 8 | 4 | 11 | 8 | 1 | 8 | 8 | 7 | 7 | 7 |
| EluB108 | 2 | 5 | 6 | 5 | 5 | 5 | 4 | 8 | 5 | 3 | 5 | 6 | 4 | 6 | 7 |
| EluBe | 5 | 3 | 5 | 3 | 3 | 4 | 3 | 4 | 3 | 2 | 2 | 4 | 5 | 5 | 3 |
| B24 | 12 | 16 | 16 | 11 | 14 | 11 | 10 | 13 | 14 | 8 | 12 | 12 | 9 | 12 | 13 |
| Eluc033 | 2 | 5 | 13 | 2 | 6 | 4 | 6 | 9 | 8 | 3 | 6 | 8 | 6 | 6 | 9 |

Population IDs are explained in Table 1 of the main document.

**B**

| loci | BAR | AWU | AMS | AMP | CHS | DON | ILZ | INN | KOS | NAB | ROT | STS | WAS | WLS | EID |
|---|---|---|---|---|---|---|---|---|---|---|---|---|---|---|---|
| Elu87 | 0 | 0 | 0 | 0 | 0 | 0 | 0 | 0 | 0 | 0 | 0 | 0 | 0 | 0 | 0 |
| Eluc045 | 0 | 0 | 0 | 0 | 0 | 0 | 0 | 0 | 0 | 0 | 0 | 0 | 0 | 0 | 0 |
| B451 | 0 | 0 | 0 | 0 | 0 | 0 | 0 | 0 | 0 | 0 | 0 | 0 | 0 | 0 | 0 |
| PkB47 | 0 | 0 | 0 | 0 | 0 | 0 | 0 | 0 | 0 | 0 | 0 | 0 | 0 | 0 | 0 |
| Elu19 | 0 | 0 | 0 | 0 | 0 | 1 | 0 | 0 | 0 | 0 | 0 | 0 | 0 | 0 | 0 |
| El02 | 0 | 1 | 0 | 0 | 0 | 0 | 0 | 0 | 0 | 0 | 0 | 0 | 0 | 0 | 0 |
| Pkb16 | 0 | 0 | 0 | 0 | 0 | 0 | 0 | 0 | 0 | 0 | 1 | 0 | 0 | 0 | 0 |
| Elu76 | 0 | 0 | 0 | 0 | 0 | 0 | 0 | 0 | 0 | 0 | 0 | 0 | 0 | 0 | 1 |
| EL27 | 0 | 0 | 0 | 0 | 0 | 0 | 0 | 0 | 0 | 0 | 0 | 0 | 0 | 0 | 0 |
| EmaD12a | 0 | 0 | 0 | 0 | 0 | 0 | 0 | 0 | 0 | 0 | 0 | 0 | 1 | 0 | 0 |
| EL01 | 0 | 0 | 0 | 0 | 0 | 0 | 0 | 0 | 0 | 0 | 0 | 0 | 0 | 0 | 0 |
| EluB108 | 0 | 0 | 0 | 0 | 0 | 0 | 0 | 0 | 0 | 0 | 0 | 0 | 0 | 0 | 0 |
| EluBe | 0 | 0 | 0 | 0 | 0 | 0 | 0 | 0 | 0 | 0 | 0 | 0 | 0 | 0 | 0 |
| B24 | 0 | 0 | 0 | 1 | 0 | 0 | 0 | 0 | 0 | 0 | 0 | 0 | 0 | 0 | 0 |
| Eluc033 | 0 | 0 | 0 | 0 | 0 | 0 | 0 | 0 | 0 | 0 | 0 | 0 | 0 | 0 | 0 |

| loci | DRS | ELB6 | ELB7 | GKB | GST | HAV | JAG | KAR | KRK | MUR | STP | EMS2 | EMS3 | NEI1 | NEI2 |
|---|---|---|---|---|---|---|---|---|---|---|---|---|---|---|---|
| Elu87 | 0 | 0 | 0 | 0 | 0 | np | 0 | 0 | 0 | 0 | 0 | 0 | 0 | 0 | 0 |
| Eluc045 | 0 | 0 | 0 | 0 | 0 | np | 0 | 0 | 0 | 0 | 0 | 0 | 0 | 0 | 0 |
| B451 | 0 | 0 | 1 | 0 | 0 | np | 0 | 0 | 0 | 0 | 1 | 2 | 0 | 0 | 0 |
| PkB47 | 0 | 0 | 0 | 0 | 0 | np | 0 | 0 | 0 | 0 | 0 | 0 | 0 | 0 | 0 |
| Elu19 | 0 | 0 | 1 | 0 | 0 | np | 0 | 0 | 0 | 0 | 0 | 0 | 0 | 0 | 0 |
| El02 | 0 | 0 | 0 | 5 | 0 | np | 0 | 0 | 0 | 2 | 0 | 0 | 0 | 0 | 0 |
| Pkb16 | 2 | 0 | 2 | 0 | 0 | np | 0 | 0 | 1 | 0 | 0 | 0 | 0 | 0 | 0 |
| Elu76 | 1 | 0 | 0 | 1 | 0 | np | 0 | 0 | 0 | 0 | 0 | 0 | 0 | 0 | 0 |
| EL27 | 0 | 0 | 0 | 0 | 0 | np | 0 | 2 | 0 | 0 | 0 | 0 | 0 | 0 | 0 |
| EmaD12a | 0 | 0 | 0 | 0 | 0 | np | 0 | 0 | 0 | 0 | 0 | 0 | 0 | 0 | 0 |
| EL01 | 0 | 0 | 0 | 0 | 0 | np | 0 | 0 | 0 | 0 | 0 | 1 | 0 | 0 | 0 |
| EluB108 | 0 | 1 | 0 | 0 | 0 | np | 0 | 0 | 0 | 0 | 0 | 0 | 0 | 0 | 0 |
| EluBe | 0 | 0 | 0 | 0 | 0 | np | 0 | 0 | 0 | 0 | 0 | 0 | 0 | 0 | 0 |
| B24 | 0 | 0 | 0 | 0 | 0 | np | 0 | 0 | 0 | 0 | 0 | 0 | 0 | 0 | 0 |
| Eluc033 | 0 | 0 | 0 | 0 | 0 | np | 0 | 0 | 0 | 0 | 0 | 0 | 0 | 0 | 0 |

| loci | NEI4 | ODE2 | ODE7 | WBS | BAS1 | BAS2 | BAS4 | BAL2 | BAL3 | BAS5 | BAL4 | BAS6 | PEE | BOS1 | BOS3 |
|---|---|---|---|---|---|---|---|---|---|---|---|---|---|---|---|
| Elu87 | np | np | 0 | 0 | 0 | 0 | 0 | 0 | 0 | 0 | 0 | 0 | 0 | 0 | 0 |
| Eluc045 | np | np | 0 | 0 | 0 | 0 | 0 | 0 | 0 | 0 | 0 | 0 | 0 | 0 | 0 |
| B451 | np | np | 0 | 0 | 0 | 0 | 2 | 0 | 1 | 0 | 0 | 0 | 0 | 0 | 0 |
| PkB47 | np | np | 2 | 0 | 0 | 2 | 0 | 2 | 0 | 0 | 0 | 0 | 0 | 0 | 0 |
| Elu19 | np | np | 0 | 0 | 0 | 0 | 0 | 0 | 0 | 0 | 0 | 0 | 0 | 0 | 0 |
| El02 | np | np | 1 | 0 | 1 | 2 | 2 | 0 | 5 | 3 | 2 | 1 | 0 | 2 | 0 |
| Pkb16 | np | np | 0 | 4 | 0 | 0 | 0 | 0 | 0 | 0 | 2 | 0 | 0 | 0 | 0 |
| Elu76 | np | np | 0 | 1 | 0 | 0 | 0 | 0 | 1 | 1 | 0 | 0 | 2 | 0 | 0 |
| EL27 | np | np | 0 | 0 | 0 | 0 | 0 | 0 | 0 | 0 | 0 | 0 | 0 | 0 | 0 |
| EmaD12a | np | np | 0 | 0 | 0 | 0 | 0 | 0 | 0 | 0 | 2 | 0 | 0 | 0 | 0 |
| EL01 | np | np | 0 | 0 | 0 | 0 | 0 | 0 | 0 | 0 | 0 | 0 | 0 | 0 | 0 |
| EluB108 | np | np | 0 | 0 | 0 | 0 | 0 | 0 | 0 | 0 | 0 | 0 | 0 | 0 | 0 |
| EluBe | np | np | 0 | 0 | 0 | 0 | 0 | 0 | 0 | 0 | 0 | 0 | 0 | 0 | 0 |
| B24 | np | np | 0 | 0 | 0 | 0 | 0 | 0 | 0 | 0 | 2 | 0 | 0 | 0 | 0 |
| Eluc033 | np | np | 0 | 0 | 0 | 0 | 0 | 0 | 0 | 0 | 0 | 0 | 0 | 0 | 0 |

| loci | BOS4 | BOS5 | MAI7 | RHE2 | GPS | HAH | KDO | MAZ | OST | SUS | WIS | GRA | UCK4 | EDS | STM |
|---|---|---|---|---|---|---|---|---|---|---|---|---|---|---|---|
| Elu87 | 0 | 0 | 1 | 0 | 1 | 0 | 0 | 0 | 0 | 0 | 0 | 0 | 0 | 0 | 0 |
| Eluc045 | 0 | 0 | 0 | 0 | 0 | 0 | 0 | 0 | 0 | 0 | 0 | 0 | 0 | 0 | 0 |
| B451 | 1 | 0 | 2 | 0 | 0 | 0 | 0 | 0 | 5 | 0 | 0 | 1 | 0 | 1 | 1 |
| PkB47 | 0 | 0 | 0 | 0 | 0 | 0 | 0 | 0 | 0 | 0 | 0 | 0 | 0 | 0 | 0 |
| Elu19 | 0 | 0 | 0 | 0 | 0 | 1 | 0 | 0 | 0 | 0 | 0 | 0 | 0 | 0 | 0 |
| El02 | 0 | 0 | 0 | 0 | 0 | 0 | 3 | 5 | 0 | 0 | 1 | 2 | 1 | 0 | 0 |
| Pkb16 | 0 | 0 | 0 | 0 | 0 | 3 | 0 | 1 | 2 | 0 | 0 | 0 | 2 | 0 | 0 |
| Elu76 | 0 | 0 | 0 | 2 | 1 | 0 | 2 | 0 | 0 | 0 | 0 | 0 | 0 | 0 | 0 |
| EL27 | 0 | 0 | 0 | 0 | 0 | 0 | 0 | 0 | 0 | 0 | 0 | 0 | 0 | 0 | 0 |
| EmaD12a | 0 | 0 | 0 | 0 | 0 | 0 | 0 | 0 | 0 | 0 | 0 | 0 | 0 | 0 | 0 |
| EL01 | 0 | 0 | 0 | 0 | 0 | 0 | 0 | 0 | 0 | 0 | 0 | 0 | 0 | 0 | 0 |
| EluB108 | 0 | 0 | 0 | 0 | 0 | 0 | 0 | 0 | 0 | 0 | 0 | 0 | 0 | 0 | 0 |
| EluBe | 0 | 0 | 0 | 0 | 0 | 0 | 0 | 0 | 0 | 0 | 0 | 0 | 0 | 0 | 0 |
| B24 | 0 | 0 | 0 | 0 | 0 | 0 | 0 | 0 | 0 | 0 | 0 | 0 | 0 | 0 | 0 |
| Eluc033 | 0 | 0 | 0 | 0 | 0 | 0 | 0 | 0 | 0 | 0 | 0 | 1 | 0 | 0 | 0 |

Population IDs are explained in Table 1 of the main document.
np = analysis not possible
