## supplemental table 2 for "Intraspecific population admixture of a top piscivore correlates with anthropogenic alteration of freshwater ecosystems"

**Table SX**: Pairwise Fst (below diagonal) and p values (above diagonal)

| | BAR | AWU | AMS | AMP | CHS | DON | ILZ | INN | KOS | NAB | ROT | STS | WAS | WLS | EID |
|---|---|---|---|---|---|---|---|---|---|---|---|---|---|---|---|
| **BAR** | | 0,0001 | 0,0001 | 0,0001 | 0,0001 | 0,0001 | 0,0001 | 0,0001 | 0,0001 | 0,0001 | 0,0001 | 0,0001 | 0,0001 | 0,0001 | 0,0001 |
| **AWU** | 0,0770 | | 0,0001 | 0,0009 | 0,0001 | 0,0002 | 0,0054 | 0,0044 | 0,0001 | 0,0020 | 0,0027 | 0,0001 | 0,0001 | 0,0001 | 0,0001 |
| **AMS** | 0,0444 | 0,0366 | | 0,0003 | 0,0001 | 0,0001 | 0,0011 | 0,0009 | 0,0010 | 0,0016 | 0,0039 | 0,0001 | 0,0001 | 0,0001 | 0,0001 |
| **AMP** | 0,0638 | 0,0634 | 0,0358 | | 0,0218 | 0,0054 | 0,1062 | 0,1804 | 0,0009 | 0,3116 | 0,0058 | 0,0001 | 0,0009 | 0,0012 | 0,0006 |
| **CHS** | 0,0464 | 0,0578 | 0,0271 | 0,0316 | | 0,0001 | 0,0355 | 0,1427 | 0,2606 | 0,5010 | 0,0941 | 0,0012 | 0,0682 | 0,0001 | 0,0039 |
| **DON** | 0,0586 | 0,0611 | 0,0415 | 0,0441 | 0,0535 | | 0,1983 | 0,0100 | 0,0003 | 0,0021 | 0,0041 | 0,0001 | 0,0001 | 0,0001 | 0,0001 |
| **ILZ** | 0,0570 | 0,0737 | 0,0465 | 0,0475 | 0,0468 | 0,0457 | | 0,4751 | 0,0167 | 0,3869 | 0,7194 | 0,0005 | 0,0024 | 0,0076 | 0,0892 |
| **INN** | 0,0675 | 0,0808 | 0,0556 | 0,0471 | 0,0467 | 0,0697 | 0,0566 | | 0,0132 | 0,3515 | 0,1310 | 0,0057 | 0,0113 | 0,0265 | 0,1115 |
| **KOS** | 0,0556 | 0,0598 | 0,0235 | 0,0375 | 0,0198 | 0,0493 | 0,0486 | 0,0576 | | 0,0378 | 0,0023 | 0,0026 | 0,0003 | 0,0003 | 0,0001 |
| **NAB** | 0,0510 | 0,0652 | 0,0357 | 0,0316 | 0,0252 | 0,0558 | 0,0449 | 0,0502 | 0,0359 | | 0,3687 | 0,0005 | 0,0020 | 0,0005 | 0,0532 |
| **ROT** | 0,0400 | 0,0590 | 0,0310 | 0,0412 | 0,0304 | 0,0504 | 0,0396 | 0,0565 | 0,0409 | 0,0326 | | 0,0001 | 0,0003 | 0,0006 | 0,0358 |
| **STS** | 0,0620 | 0,0527 | 0,0187 | 0,0367 | 0,0238 | 0,0473 | 0,0519 | 0,0491 | 0,0217 | 0,0389 | 0,0422 | | 0,0002 | 0,0001 | 0,0001 |
| **WAS** | 0,0564 | 0,0688 | 0,0277 | 0,0307 | 0,0211 | 0,0551 | 0,0530 | 0,0482 | 0,0283 | 0,0360 | 0,0374 | 0,0204 | | 0,0001 | 0,0001 |
| **WLS** | 0,0663 | 0,0687 | 0,0332 | 0,0297 | 0,0283 | 0,0443 | 0,0441 | 0,0458 | 0,0271 | 0,0395 | 0,0385 | 0,0252 | 0,0284 | | 0,0001 |
| **EID** | 0,0450 | 0,0664 | 0,0376 | 0,0359 | 0,0235 | 0,0594 | 0,0357 | 0,0399 | 0,0401 | 0,0270 | 0,0280 | 0,0476 | 0,0350 | 0,0426 | |
| **DRS** | 0,0633 | 0,0742 | 0,0529 | 0,0380 | 0,0351 | 0,0644 | 0,0532 | 0,0406 | 0,0490 | 0,0418 | 0,0454 | 0,0555 | 0,0403 | 0,0440 | 0,0315 |
| **ELB6** | 0,0525 | 0,0835 | 0,0522 | 0,0461 | 0,0412 | 0,0733 | 0,0397 | 0,0573 | 0,0534 | 0,0369 | 0,0386 | 0,0578 | 0,0392 | 0,0555 | 0,0228 |
| **ELB7** | 0,0464 | 0,0751 | 0,0483 | 0,0343 | 0,0415 | 0,0556 | 0,0385 | 0,0461 | 0,0576 | 0,0353 | 0,0361 | 0,0592 | 0,0434 | 0,0525 | 0,0217 |
| **GKB** | 0,0574 | 0,0844 | 0,0569 | 0,0409 | 0,0456 | 0,0628 | 0,0507 | 0,0642 | 0,0644 | 0,0368 | 0,0367 | 0,0687 | 0,0514 | 0,0552 | 0,0357 |
| **GST** | 0,0575 | 0,0667 | 0,0470 | 0,0346 | 0,0385 | 0,0503 | 0,0479 | 0,0518 | 0,0444 | 0,0398 | 0,0448 | 0,0546 | 0,0420 | 0,0499 | 0,0339 |
| **HAV** | 0,0760 | 0,0747 | 0,0594 | 0,0653 | 0,0601 | 0,0717 | 0,0583 | 0,0862 | 0,0712 | 0,0543 | 0,0537 | 0,0786 | 0,0654 | 0,0802 | 0,0392 |
| **JAG** | 0,0497 | 0,0702 | 0,0459 | 0,0399 | 0,0380 | 0,0499 | 0,0454 | 0,0440 | 0,0505 | 0,0376 | 0,0373 | 0,0521 | 0,0372 | 0,0434 | 0,0289 |
| **KAR** | 0,0425 | 0,0652 | 0,0411 | 0,0276 | 0,0277 | 0,0451 | 0,0365 | 0,0377 | 0,0412 | 0,0278 | 0,0297 | 0,0484 | 0,0331 | 0,0390 | 0,0170 |
| **KRK** | 0,0476 | 0,0817 | 0,0502 | 0,0447 | 0,0393 | 0,0679 | 0,0446 | 0,0474 | 0,0560 | 0,0393 | 0,0400 | 0,0557 | 0,0371 | 0,0546 | 0,0249 |
| **MUR** | 0,0811 | 0,0931 | 0,0753 | 0,0606 | 0,0656 | 0,0750 | 0,0669 | 0,0664 | 0,0806 | 0,0661 | 0,0705 | 0,0794 | 0,0668 | 0,0750 | 0,0591 |
| **STP** | 0,0581 | 0,0926 | 0,0593 | 0,0551 | 0,0401 | 0,0675 | 0,0428 | 0,0537 | 0,0556 | 0,0397 | 0,0499 | 0,0640 | 0,0534 | 0,0577 | 0,0324 |
| **EMS2** | 0,0534 | 0,0739 | 0,0552 | 0,0478 | 0,0411 | 0,0732 | 0,0472 | 0,0551 | 0,0618 | 0,0325 | 0,0393 | 0,0629 | 0,0540 | 0,0617 | 0,0246 |
| **EMS3** | 0,0404 | 0,0814 | 0,0492 | 0,0491 | 0,0397 | 0,0695 | 0,0521 | 0,0650 | 0,0487 | 0,0470 | 0,0477 | 0,0572 | 0,0410 | 0,0610 | 0,0367 |
| **NEI1** | 0,0632 | 0,0937 | 0,0694 | 0,0566 | 0,0720 | 0,0663 | 0,0710 | 0,0883 | 0,0756 | 0,0599 | 0,0564 | 0,0884 | 0,0722 | 0,0752 | 0,0534 |
| **NEI2** | 0,0675 | 0,1009 | 0,0730 | 0,0635 | 0,0655 | 0,0765 | 0,0695 | 0,0908 | 0,0761 | 0,0624 | 0,0525 | 0,0848 | 0,0693 | 0,0695 | 0,0528 |
| **NEI4** | 0,0733 | 0,1062 | 0,0627 | 0,0541 | 0,0591 | 0,0823 | 0,0793 | 0,0914 | 0,0613 | 0,0576 | 0,0581 | 0,0688 | 0,0597 | 0,0690 | 0,0592 |
| **ODE2** | 0,0422 | 0,0967 | 0,0659 | 0,0699 | 0,0561 | 0,0612 | 0,0634 | 0,0796 | 0,0727 | 0,0588 | 0,0461 | 0,0826 | 0,0734 | 0,0814 | 0,0478 |
| **ODE7** | 0,0396 | 0,0770 | 0,0422 | 0,0314 | 0,0359 | 0,0437 | 0,0416 | 0,0418 | 0,0441 | 0,0333 | 0,0342 | 0,0489 | 0,0363 | 0,0411 | 0,0245 |
| **WBS** | 0,0484 | 0,0896 | 0,0549 | 0,0541 | 0,0499 | 0,0552 | 0,0604 | 0,0687 | 0,0548 | 0,0459 | 0,0477 | 0,0674 | 0,0626 | 0,0667 | 0,0447 |
| **BAL2** | 0,0358 | 0,0939 | 0,0596 | 0,0722 | 0,0648 | 0,0606 | 0,0604 | 0,0740 | 0,0670 | 0,0693 | 0,0559 | 0,0783 | 0,0689 | 0,0761 | 0,0500 |
| **BAL3** | 0,0462 | 0,0901 | 0,0548 | 0,0687 | 0,0506 | 0,0808 | 0,0651 | 0,0784 | 0,0668 | 0,0564 | 0,0567 | 0,0711 | 0,0653 | 0,0742 | 0,0398 |
| **BAL4** | 0,0358 | 0,0773 | 0,0440 | 0,0491 | 0,0411 | 0,0561 | 0,0431 | 0,0554 | 0,0542 | 0,0391 | 0,0382 | 0,0587 | 0,0466 | 0,0553 | 0,0253 |
| **PEE** | 0,0285 | 0,0740 | 0,0420 | 0,0416 | 0,0405 | 0,0471 | 0,0482 | 0,0540 | 0,0509 | 0,0381 | 0,0334 | 0,0584 | 0,0441 | 0,0541 | 0,0289 |
| **BOS1** | 0,0523 | 0,0565 | 0,0304 | 0,0469 | 0,0424 | 0,0546 | 0,0467 | 0,0630 | 0,0402 | 0,0436 | 0,0320 | 0,0425 | 0,0401 | 0,0422 | 0,0436 |
| **BOS3** | 0,0730 | 0,0674 | 0,0445 | 0,0617 | 0,0590 | 0,0751 | 0,0586 | 0,0814 | 0,0579 | 0,0573 | 0,0503 | 0,0538 | 0,0556 | 0,0536 | 0,0621 |
| **BOS4** | 0,0699 | 0,1014 | 0,0653 | 0,0854 | 0,0712 | 0,0993 | 0,0785 | 0,0935 | 0,0780 | 0,0772 | 0,0668 | 0,0705 | 0,0611 | 0,0790 | 0,0722 |
| **BOS5** | 0,0430 | 0,0791 | 0,0435 | 0,0596 | 0,0442 | 0,0647 | 0,0558 | 0,0784 | 0,0519 | 0,0523 | 0,0427 | 0,0540 | 0,0501 | 0,0609 | 0,0483 |
| **MAI7** | 0,0299 | 0,0450 | 0,0102 | 0,0291 | 0,0191 | 0,0381 | 0,0351 | 0,0451 | 0,0216 | 0,0244 | 0,0220 | 0,0227 | 0,0226 | 0,0301 | 0,0213 |
| **RHE2** | 0,0752 | 0,1240 | 0,0775 | 0,0862 | 0,0666 | 0,1067 | 0,0791 | 0,0964 | 0,0837 | 0,0721 | 0,0789 | 0,0884 | 0,0804 | 0,0856 | 0,0543 |
| **GPS** | 0,0523 | 0,0556 | 0,0290 | 0,0371 | 0,0288 | 0,0562 | 0,0541 | 0,0550 | 0,0338 | 0,0341 | 0,0292 | 0,0384 | 0,0287 | 0,0412 | 0,0285 |
| **HAH** | 0,0487 | 0,0962 | 0,0646 | 0,0603 | 0,0489 | 0,0652 | 0,0552 | 0,0676 | 0,0618 | 0,0517 | 0,0519 | 0,0728 | 0,0641 | 0,0742 | 0,0421 |
| **KDO** | 0,0972 | 0,1322 | 0,0858 | 0,0718 | 0,0716 | 0,0969 | 0,0873 | 0,0967 | 0,0828 | 0,0734 | 0,0862 | 0,0893 | 0,0628 | 0,0893 | 0,0634 |
| **SUS** | 0,1037 | 0,1281 | 0,0972 | 0,0902 | 0,0819 | 0,1132 | 0,0924 | 0,1119 | 0,0878 | 0,1031 | 0,0997 | 0,1010 | 0,0852 | 0,1009 | 0,0823 |
| **WIS** | 0,0568 | 0,0703 | 0,0353 | 0,0330 | 0,0294 | 0,0623 | 0,0551 | 0,0598 | 0,0343 | 0,0300 | 0,0385 | 0,0361 | 0,0227 | 0,0450 | 0,0340 |
| **GRA** | 0,0527 | 0,0879 | 0,0500 | 0,0462 | 0,0415 | 0,0673 | 0,0595 | 0,0692 | 0,0565 | 0,0424 | 0,0497 | 0,0588 | 0,0478 | 0,0612 | 0,0374 |
| **UCK4** | 0,0531 | 0,0739 | 0,0470 | 0,0483 | 0,0498 | 0,0536 | 0,0590 | 0,0588 | 0,0636 | 0,0482 | 0,0439 | 0,0523 | 0,0476 | 0,0646 | 0,0430 |
| **EDS** | 0,0372 | 0,0626 | 0,0342 | 0,0288 | 0,0278 | 0,0537 | 0,0508 | 0,0427 | 0,0408 | 0,0304 | 0,0327 | 0,0369 | 0,0270 | 0,0367 | 0,0323 |
| **STM** | 0,0453 | 0,0742 | 0,0539 | 0,0475 | 0,0435 | 0,0679 | 0,0470 | 0,0615 | 0,0616 | 0,0362 | 0,0332 | 0,0636 | 0,0514 | 0,0629 | 0,0266 |

| DRS | ELB6 | ELB7 | GKB | GST | HAV | JAG | KAR | KRK | MUR | STP | EMS2 | EMS3 | NEI1 | NEI2 | NEI4 |
|---|---|---|---|---|---|---|---|---|---|---|---|---|---|---|---|
| 0,0001 | 0,0001 | 0,0001 | 0,0001 | 0,0001 | 0,0001 | 0,0001 | 0,0001 | 0,0001 | 0,0001 | 0,0001 | 0,0001 | 0,0001 | 0,0001 | 0,0001 | 0,0013 |
| 0,0001 | 0,0001 | 0,0001 | 0,0001 | 0,0001 | 0,0001 | 0,0001 | 0,0001 | 0,0001 | 0,0001 | 0,0001 | 0,0001 | 0,0001 | 0,0002 | 0,0018 | 0,0033 |
| 0,0001 | 0,0001 | 0,0001 | 0,0001 | 0,0001 | 0,0001 | 0,0001 | 0,0001 | 0,0001 | 0,0001 | 0,0001 | 0,0001 | 0,0001 | 0,0001 | 0,0001 | 0,0065 |
| 0,0006 | 0,0001 | 0,0010 | 0,0003 | 0,0001 | 0,0001 | 0,0001 | 0,0015 | 0,0001 | 0,0001 | 0,0001 | 0,0001 | 0,0002 | 0,0006 | 0,0078 | 0,2403 |
| 0,0001 | 0,0001 | 0,0001 | 0,0001 | 0,0001 | 0,0001 | 0,0001 | 0,0001 | 0,0001 | 0,0001 | 0,0001 | 0,0001 | 0,0005 | 0,0001 | 0,0019 | 0,0383 |
| 0,0001 | 0,0001 | 0,0001 | 0,0001 | 0,0001 | 0,0001 | 0,0003 | 0,0001 | 0,0001 | 0,0001 | 0,0001 | 0,0001 | 0,0001 | 0,0022 | 0,0027 | 0,0158 |
| 0,0062 | 0,0513 | 0,1766 | 0,0104 | 0,0011 | 0,0350 | 0,0235 | 0,0581 | 0,0036 | 0,0001 | 0,0545 | 0,0202 | 0,0439 | 0,0106 | 0,1761 | 0,1943 |
| 0,1852 | 0,0030 | 0,1295 | 0,0027 | 0,0045 | 0,0072 | 0,0615 | 0,1941 | 0,0123 | 0,0006 | 0,0406 | 0,0287 | 0,0096 | 0,0021 | 0,0118 | 0,0812 |
| 0,0001 | 0,0001 | 0,0001 | 0,0001 | 0,0001 | 0,0001 | 0,0001 | 0,0001 | 0,0001 | 0,0001 | 0,0001 | 0,0001 | 0,0001 | 0,0001 | 0,0005 | 0,0036 |
| 0,0042 | 0,0020 | 0,0231 | 0,0133 | 0,0015 | 0,0005 | 0,0080 | 0,0503 | 0,0001 | 0,0001 | 0,0151 | 0,0555 | 0,0106 | 0,0129 | 0,0390 | 0,3057 |
| 0,0001 | 0,0003 | 0,0073 | 0,0027 | 0,0001 | 0,0019 | 0,0036 | 0,0100 | 0,0001 | 0,0001 | 0,0010 | 0,0028 | 0,0025 | 0,0066 | 0,1777 | 0,2734 |
| 0,0001 | 0,0001 | 0,0001 | 0,0001 | 0,0001 | 0,0001 | 0,0001 | 0,0001 | 0,0001 | 0,0001 | 0,0001 | 0,0001 | 0,0001 | 0,0001 | 0,0001 | 0,0001 |
| 0,0001 | 0,0001 | 0,0001 | 0,0001 | 0,0001 | 0,0001 | 0,0001 | 0,0001 | 0,0001 | 0,0001 | 0,0001 | 0,0001 | 0,0001 | 0,0001 | 0,0001 | 0,0036 |
| 0,0001 | 0,0001 | 0,0001 | 0,0001 | 0,0001 | 0,0001 | 0,0001 | 0,0001 | 0,0001 | 0,0001 | 0,0001 | 0,0001 | 0,0001 | 0,0001 | 0,0002 | 0,0028 |
| 0,0001 | 0,0001 | 0,0062 | 0,0001 | 0,0001 | 0,0001 | 0,0001 | 0,0007 | 0,0001 | 0,0001 | 0,0001 | 0,0001 | 0,0001 | 0,0010 | 0,0058 | 0,0194 |
| | 0,0001 | 0,0001 | 0,0001 | 0,0001 | 0,0001 | 0,0001 | 0,0003 | 0,0001 | 0,0001 | 0,0001 | 0,0001 | 0,0001 | 0,0001 | 0,0013 | 0,0076 |
| 0,0423 | | 0,0014 | 0,0001 | 0,0001 | 0,0001 | 0,0001 | 0,0001 | 0,0001 | 0,0001 | 0,0001 | 0,0001 | 0,0019 | 0,0001 | 0,0005 | 0,0033 |
| 0,0383 | 0,0238 | | 0,0001 | 0,0001 | 0,0001 | 0,0032 | 0,0922 | 0,0001 | 0,0001 | 0,0001 | 0,0001 | 0,0001 | 0,0038 | 0,0714 | 0,0089 |
| 0,0384 | 0,0356 | 0,0245 | | 0,0001 | 0,0001 | 0,0001 | 0,0001 | 0,0001 | 0,0001 | 0,0001 | 0,0001 | 0,0001 | 0,0109 | 0,0105 | 0,0593 |
| 0,0287 | 0,0450 | 0,0283 | 0,0384 | | 0,0001 | 0,0001 | 0,0001 | 0,0001 | 0,0001 | 0,0001 | 0,0001 | 0,0001 | 0,0001 | 0,0006 | 0,0016 |
| 0,0550 | 0,0465 | 0,0412 | 0,0460 | 0,0410 | | 0,0004 | 0,0001 | 0,0001 | 0,0001 | 0,0001 | 0,0001 | 0,0001 | 0,0124 | 0,0043 | 0,0029 |
| 0,0327 | 0,0362 | 0,0255 | 0,0339 | 0,0267 | 0,0433 | | 0,4080 | 0,0002 | 0,0001 | 0,0001 | 0,0001 | 0,0004 | 0,0005 | 0,0068 | 0,0017 |
| 0,0210 | 0,0245 | 0,0158 | 0,0236 | 0,0182 | 0,0390 | 0,0142 | | 0,0001 | 0,0001 | 0,0001 | 0,0001 | 0,0019 | 0,0004 | 0,0025 | 0,0020 |
| 0,0356 | 0,0174 | 0,0227 | 0,0336 | 0,0355 | 0,0469 | 0,0241 | 0,0214 | | 0,0001 | 0,0001 | 0,0001 | 0,0198 | 0,0001 | 0,0001 | 0,0002 |
| 0,0365 | 0,0612 | 0,0504 | 0,0545 | 0,0349 | 0,0553 | 0,0404 | 0,0368 | 0,0441 | | 0,0001 | 0,0001 | 0,0001 | 0,0001 | 0,0001 | 0,0005 |
| 0,0451 | 0,0346 | 0,0334 | 0,0465 | 0,0445 | 0,0637 | 0,0391 | 0,0303 | 0,0304 | 0,0632 | | 0,0001 | 0,0001 | 0,0001 | 0,0003 | 0,0052 |
| 0,0397 | 0,0294 | 0,0318 | 0,0344 | 0,0427 | 0,0405 | 0,0404 | 0,0279 | 0,0295 | 0,0526 | 0,0434 | | 0,0002 | 0,0001 | 0,0019 | 0,0016 |
| 0,0508 | 0,0305 | 0,0392 | 0,0484 | 0,0436 | 0,0556 | 0,0370 | 0,0310 | 0,0224 | 0,0554 | 0,0453 | 0,0423 | | 0,0001 | 0,0005 | 0,0242 |
| 0,0723 | 0,0634 | 0,0471 | 0,0464 | 0,0567 | 0,0659 | 0,0536 | 0,0464 | 0,0663 | 0,0956 | 0,0793 | 0,0692 | 0,0680 | | 0,4871 | 0,3439 |
| 0,0711 | 0,0627 | 0,0475 | 0,0530 | 0,0587 | 0,0737 | 0,0556 | 0,0511 | 0,0617 | 0,0854 | 0,0775 | 0,0624 | 0,0734 | 0,0451 | | 0,1309 |
| 0,0791 | 0,0639 | 0,0689 | 0,0591 | 0,0746 | 0,0887 | 0,0770 | 0,0646 | 0,0716 | 0,1158 | 0,0804 | 0,0783 | 0,0664 | 0,0647 | 0,0804 | |
| 0,0643 | 0,0592 | 0,0437 | 0,0476 | 0,0553 | 0,0917 | 0,0470 | 0,0353 | 0,0546 | 0,0776 | 0,0543 | 0,0616 | 0,0564 | 0,0609 | 0,0705 | 0,0835 |
| 0,0379 | 0,0317 | 0,0217 | 0,0306 | 0,0321 | 0,0546 | 0,0258 | 0,0179 | 0,0285 | 0,0579 | 0,0345 | 0,0406 | 0,0358 | 0,0392 | 0,0464 | 0,0502 |
| 0,0605 | 0,0583 | 0,0490 | 0,0416 | 0,0534 | 0,0682 | 0,0544 | 0,0408 | 0,0585 | 0,0807 | 0,0613 | 0,0582 | 0,0572 | 0,0457 | 0,0613 | 0,0513 |
| 0,0714 | 0,0581 | 0,0464 | 0,0593 | 0,0538 | 0,0765 | 0,0479 | 0,0441 | 0,0517 | 0,0780 | 0,0652 | 0,0671 | 0,0535 | 0,0585 | 0,0631 | 0,0874 |
| 0,0713 | 0,0465 | 0,0570 | 0,0613 | 0,0693 | 0,0709 | 0,0585 | 0,0532 | 0,0500 | 0,0892 | 0,0546 | 0,0497 | 0,0405 | 0,0722 | 0,0699 | 0,0711 |
| 0,0488 | 0,0290 | 0,0286 | 0,0303 | 0,0436 | 0,0480 | 0,0330 | 0,0280 | 0,0274 | 0,0573 | 0,0402 | 0,0378 | 0,0313 | 0,0469 | 0,0535 | 0,0543 |
| 0,0447 | 0,0387 | 0,0248 | 0,0324 | 0,0348 | 0,0544 | 0,0314 | 0,0224 | 0,0352 | 0,0619 | 0,0444 | 0,0430 | 0,0400 | 0,0330 | 0,0445 | 0,0549 |
| 0,0512 | 0,0472 | 0,0519 | 0,0627 | 0,0582 | 0,0716 | 0,0511 | 0,0401 | 0,0528 | 0,0830 | 0,0604 | 0,0532 | 0,0539 | 0,0753 | 0,0742 | 0,0804 |
| 0,0653 | 0,0634 | 0,0724 | 0,0817 | 0,0803 | 0,0850 | 0,0654 | 0,0628 | 0,0706 | 0,0993 | 0,0767 | 0,0694 | 0,0685 | 0,0967 | 0,0933 | 0,0971 |
| 0,0795 | 0,0566 | 0,0829 | 0,0917 | 0,0957 | 0,1016 | 0,0722 | 0,0726 | 0,0632 | 0,1095 | 0,0751 | 0,0732 | 0,0537 | 0,1228 | 0,1223 | 0,1101 |
| 0,0578 | 0,0460 | 0,0584 | 0,0626 | 0,0636 | 0,0711 | 0,0611 | 0,0484 | 0,0505 | 0,0809 | 0,0561 | 0,0549 | 0,0393 | 0,0826 | 0,0851 | 0,0746 |
| 0,0396 | 0,0301 | 0,0331 | 0,0397 | 0,0362 | 0,0440 | 0,0344 | 0,0257 | 0,0325 | 0,0635 | 0,0410 | 0,0358 | 0,0313 | 0,0551 | 0,0601 | 0,0447 |
| 0,0810 | 0,0559 | 0,0737 | 0,0733 | 0,0881 | 0,0864 | 0,0719 | 0,0694 | 0,0632 | 0,1061 | 0,0641 | 0,0636 | 0,0569 | 0,0891 | 0,0857 | 0,0884 |
| 0,0523 | 0,0414 | 0,0445 | 0,0515 | 0,0447 | 0,0583 | 0,0381 | 0,0317 | 0,0404 | 0,0763 | 0,0605 | 0,0501 | 0,0406 | 0,0587 | 0,0684 | 0,0586 |
| 0,0533 | 0,0522 | 0,0423 | 0,0458 | 0,0445 | 0,0705 | 0,0501 | 0,0330 | 0,0454 | 0,0638 | 0,0455 | 0,0510 | 0,0537 | 0,0671 | 0,0683 | 0,0732 |
| 0,0836 | 0,0603 | 0,0639 | 0,0657 | 0,0638 | 0,0782 | 0,0596 | 0,0571 | 0,0529 | 0,0881 | 0,0670 | 0,0782 | 0,0530 | 0,0855 | 0,1000 | 0,0760 |
| 0,0738 | 0,0804 | 0,0819 | 0,0866 | 0,0661 | 0,0845 | 0,0820 | 0,0705 | 0,0744 | 0,0790 | 0,0930 | 0,0899 | 0,0735 | 0,1135 | 0,1126 | 0,1209 |
| 0,0506 | 0,0372 | 0,0431 | 0,0445 | 0,0398 | 0,0524 | 0,0402 | 0,0353 | 0,0379 | 0,0741 | 0,0494 | 0,0454 | 0,0383 | 0,0637 | 0,0762 | 0,0434 |
| 0,0611 | 0,0380 | 0,0391 | 0,0411 | 0,0493 | 0,0614 | 0,0445 | 0,0393 | 0,0339 | 0,0754 | 0,0437 | 0,0491 | 0,0338 | 0,0653 | 0,0689 | 0,0492 |
| 0,0596 | 0,0487 | 0,0412 | 0,0448 | 0,0457 | 0,0581 | 0,0426 | 0,0367 | 0,0410 | 0,0569 | 0,0548 | 0,0501 | 0,0497 | 0,0691 | 0,0767 | 0,0658 |
| 0,0385 | 0,0364 | 0,0322 | 0,0328 | 0,0377 | 0,0566 | 0,0324 | 0,0257 | 0,0322 | 0,0590 | 0,0452 | 0,0306 | 0,0330 | 0,0577 | 0,0649 | 0,0569 |
| 0,0369 | 0,0287 | 0,0301 | 0,0295 | 0,0373 | 0,0336 | 0,0343 | 0,0244 | 0,0267 | 0,0449 | 0,0457 | 0,0125 | 0,0339 | 0,0623 | 0,0570 | 0,0711 |

| ODE2 | ODE7 | WBS | BAL2 | BAL3 | BAL4 | PEE | BOS1 | BOS3 | BOS4 | BOS5 | MAI7 | RHE2 | GPS | HAH | KDO |
|---|---|---|---|---|---|---|---|---|---|---|---|---|---|---|---|
| 0,0001 | 0,0001 | 0,0001 | 0,0001 | 0,0001 | 0,0001 | 0,0001 | 0,0001 | 0,0001 | 0,0001 | 0,0001 | 0,0001 | 0,0001 | 0,0001 | 0,0001 | 0,0001 |
| 0,0001 | 0,0001 | 0,0001 | 0,0001 | 0,0001 | 0,0001 | 0,0001 | 0,0001 | 0,0001 | 0,0001 | 0,0001 | 0,0001 | 0,0001 | 0,0001 | 0,0001 | 0,0001 |
| 0,0001 | 0,0001 | 0,0001 | 0,0001 | 0,0001 | 0,0001 | 0,0001 | 0,0001 | 0,0001 | 0,0001 | 0,0001 | 0,1728 | 0,0001 | 0,0001 | 0,0001 | 0,0001 |
| 0,0001 | 0,0005 | 0,0001 | 0,0001 | 0,0001 | 0,0001 | 0,0001 | 0,0001 | 0,0001 | 0,0001 | 0,0001 | 0,0058 | 0,0001 | 0,0001 | 0,0001 | 0,0001 |
| 0,0001 | 0,0001 | 0,0001 | 0,0001 | 0,0001 | 0,0001 | 0,0001 | 0,0001 | 0,0001 | 0,0001 | 0,0001 | 0,0609 | 0,0001 | 0,0001 | 0,0001 | 0,0001 |
| 0,0002 | 0,0001 | 0,0001 | 0,0001 | 0,0001 | 0,0001 | 0,0001 | 0,0001 | 0,0001 | 0,0001 | 0,0001 | 0,0001 | 0,0001 | 0,0001 | 0,0001 | 0,0001 |
| 0,0293 | 0,0127 | 0,0001 | 0,0001 | 0,0001 | 0,0331 | 0,0018 | 0,0015 | 0,0018 | 0,0001 | 0,0006 | 0,1031 | 0,0001 | 0,0013 | 0,0005 | 0,0001 |
| 0,0223 | 0,0629 | 0,0001 | 0,0001 | 0,0001 | 0,0056 | 0,0015 | 0,0001 | 0,0002 | 0,0002 | 0,0001 | 0,0359 | 0,0002 | 0,0042 | 0,0001 | 0,0001 |
| 0,0001 | 0,0001 | 0,0001 | 0,0001 | 0,0001 | 0,0001 | 0,0001 | 0,0001 | 0,0001 | 0,0001 | 0,0001 | 0,0116 | 0,0001 | 0,0001 | 0,0001 | 0,0001 |
| 0,0036 | 0,0054 | 0,0001 | 0,0001 | 0,0001 | 0,0013 | 0,0005 | 0,0002 | 0,0004 | 0,0001 | 0,0002 | 0,2018 | 0,0001 | 0,0059 | 0,0001 | 0,0001 |
| 0,0403 | 0,0008 | 0,0001 | 0,0001 | 0,0001 | 0,0002 | 0,0010 | 0,0018 | 0,0010 | 0,0001 | 0,0006 | 0,2605 | 0,0001 | 0,0221 | 0,0001 | 0,0001 |
| 0,0001 | 0,0001 | 0,0001 | 0,0001 | 0,0001 | 0,0001 | 0,0001 | 0,0001 | 0,0001 | 0,0001 | 0,0001 | 0,0001 | 0,0001 | 0,0001 | 0,0001 | 0,0001 |
| 0,0001 | 0,0001 | 0,0001 | 0,0001 | 0,0001 | 0,0001 | 0,0001 | 0,0001 | 0,0001 | 0,0001 | 0,0001 | 0,0001 | 0,0001 | 0,0001 | 0,0001 | 0,0001 |
| 0,0001 | 0,0001 | 0,0001 | 0,0001 | 0,0001 | 0,0001 | 0,0001 | 0,0001 | 0,0001 | 0,0001 | 0,0001 | 0,0001 | 0,0001 | 0,0001 | 0,0001 | 0,0001 |
| 0,0001 | 0,0001 | 0,0001 | 0,0001 | 0,0001 | 0,0001 | 0,0001 | 0,0001 | 0,0001 | 0,0001 | 0,0001 | 0,0001 | 0,0001 | 0,0001 | 0,0001 | 0,0001 |
| 0,0001 | 0,0001 | 0,0001 | 0,0001 | 0,0001 | 0,0001 | 0,0001 | 0,0001 | 0,0001 | 0,0001 | 0,0001 | 0,0001 | 0,0001 | 0,0001 | 0,0001 | 0,0001 |
| 0,0001 | 0,0001 | 0,0001 | 0,0001 | 0,0001 | 0,0001 | 0,0001 | 0,0001 | 0,0001 | 0,0001 | 0,0001 | 0,0001 | 0,0001 | 0,0001 | 0,0001 | 0,0001 |
| 0,0002 | 0,0009 | 0,0001 | 0,0001 | 0,0001 | 0,0001 | 0,0003 | 0,0001 | 0,0001 | 0,0001 | 0,0001 | 0,0001 | 0,0001 | 0,0001 | 0,0001 | 0,0001 |
| 0,0001 | 0,0001 | 0,0001 | 0,0001 | 0,0001 | 0,0001 | 0,0001 | 0,0001 | 0,0001 | 0,0001 | 0,0001 | 0,0001 | 0,0001 | 0,0001 | 0,0001 | 0,0001 |
| 0,0001 | 0,0001 | 0,0001 | 0,0001 | 0,0001 | 0,0001 | 0,0001 | 0,0001 | 0,0001 | 0,0001 | 0,0001 | 0,0001 | 0,0001 | 0,0001 | 0,0001 | 0,0001 |
| 0,0001 | 0,0001 | 0,0001 | 0,0001 | 0,0001 | 0,0001 | 0,0001 | 0,0001 | 0,0001 | 0,0001 | 0,0001 | 0,0001 | 0,0001 | 0,0001 | 0,0001 | 0,0001 |
| 0,0004 | 0,0001 | 0,0001 | 0,0001 | 0,0001 | 0,0001 | 0,0001 | 0,0001 | 0,0001 | 0,0001 | 0,0001 | 0,0001 | 0,0001 | 0,0001 | 0,0001 | 0,0001 |
| 0,0013 | 0,0001 | 0,0001 | 0,0001 | 0,0001 | 0,0001 | 0,0001 | 0,0001 | 0,0001 | 0,0001 | 0,0001 | 0,0001 | 0,0001 | 0,0001 | 0,0001 | 0,0001 |
| 0,0001 | 0,0001 | 0,0001 | 0,0001 | 0,0001 | 0,0001 | 0,0001 | 0,0001 | 0,0001 | 0,0001 | 0,0001 | 0,0001 | 0,0001 | 0,0001 | 0,0001 | 0,0001 |
| 0,0001 | 0,0001 | 0,0001 | 0,0001 | 0,0001 | 0,0001 | 0,0001 | 0,0001 | 0,0001 | 0,0001 | 0,0001 | 0,0001 | 0,0001 | 0,0001 | 0,0001 | 0,0001 |
| 0,0001 | 0,0001 | 0,0001 | 0,0001 | 0,0001 | 0,0001 | 0,0001 | 0,0001 | 0,0001 | 0,0001 | 0,0001 | 0,0001 | 0,0001 | 0,0001 | 0,0001 | 0,0001 |
| 0,0001 | 0,0001 | 0,0001 | 0,0001 | 0,0001 | 0,0001 | 0,0001 | 0,0001 | 0,0001 | 0,0001 | 0,0001 | 0,0001 | 0,0001 | 0,0001 | 0,0001 | 0,0001 |
| 0,0004 | 0,0001 | 0,0001 | 0,0001 | 0,0001 | 0,0023 | 0,0001 | 0,0001 | 0,0001 | 0,0001 | 0,0008 | 0,0008 | 0,0001 | 0,0001 | 0,0001 | 0,0001 |
| 0,1134 | 0,0089 | 0,0035 | 0,0001 | 0,0001 | 0,0028 | 0,1393 | 0,0001 | 0,0001 | 0,0001 | 0,0001 | 0,0001 | 0,0001 | 0,0001 | 0,0001 | 0,0001 |
| 0,0235 | 0,0075 | 0,0002 | 0,0004 | 0,0001 | 0,0093 | 0,0349 | 0,0001 | 0,0005 | 0,0002 | 0,0001 | 0,0007 | 0,0001 | 0,0001 | 0,0001 | 0,0001 |
| 0,0140 | 0,0880 | 0,1108 | 0,0001 | 0,0002 | 0,0854 | 0,0455 | 0,0001 | 0,0006 | 0,0014 | 0,0013 | 0,2415 | 0,0006 | 0,0114 | 0,0033 | 0,0002 |
|  | 0,0268 | 0,0001 | 0,0001 | 0,0001 | 0,0001 | 0,2593 | 0,0001 | 0,0001 | 0,0001 | 0,0001 | 0,0001 | 0,0001 | 0,0001 | 0,0001 | 0,0001 |
| 0,0349 |  | 0,0001 | 0,0001 | 0,0001 | 0,0001 | 0,0001 | 0,0001 | 0,0001 | 0,0001 | 0,0001 | 0,0001 | 0,0001 | 0,0001 | 0,0001 | 0,0001 |
| 0,0439 | 0,0324 |  | 0,0001 | 0,0001 | 0,0001 | 0,0001 | 0,0001 | 0,0001 | 0,0001 | 0,0001 | 0,0001 | 0,0001 | 0,0001 | 0,0001 | 0,0001 |
| 0,0456 | 0,0347 | 0,0439 |  | 0,0001 | 0,0001 | 0,0001 | 0,0001 | 0,0001 | 0,0001 | 0,0001 | 0,0001 | 0,0001 | 0,0001 | 0,0001 | 0,0001 |
| 0,0740 | 0,0506 | 0,0600 | 0,0614 |  | 0,0001 | 0,0001 | 0,0001 | 0,0001 | 0,0001 | 0,0001 | 0,0001 | 0,0029 | 0,0001 | 0,0001 | 0,0001 |
| 0,0408 | 0,0223 | 0,0330 | 0,0273 | 0,0326 |  | 0,0001 | 0,0001 | 0,0001 | 0,0001 | 0,0001 | 0,0001 | 0,0001 | 0,0001 | 0,0001 | 0,0001 |
| 0,0273 | 0,0183 | 0,0240 | 0,0270 | 0,0477 | 0,0200 |  | 0,0001 | 0,0001 | 0,0001 | 0,0001 | 0,0001 | 0,0001 | 0,0001 | 0,0001 | 0,0001 |
| 0,0667 | 0,0478 | 0,0676 | 0,0684 | 0,0677 | 0,0538 | 0,0496 |  | 0,2815 | 0,0001 | 0,0001 | 0,0001 | 0,0001 | 0,0001 | 0,0001 | 0,0001 |
| 0,1012 | 0,0704 | 0,0965 | 0,1009 | 0,0798 | 0,0777 | 0,0772 | 0,0184 |  | 0,0001 | 0,0001 | 0,0001 | 0,0001 | 0,0001 | 0,0001 | 0,0001 |
| 0,1055 | 0,0781 | 0,1114 | 0,1018 | 0,0692 | 0,0771 | 0,0880 | 0,0443 | 0,0472 |  | 0,0241 | 0,0001 | 0,0001 | 0,0001 | 0,0001 | 0,0001 |
| 0,0672 | 0,0562 | 0,0656 | 0,0766 | 0,0487 | 0,0538 | 0,0572 | 0,0291 | 0,0443 | 0,0367 |  | 0,0001 | 0,0001 | 0,0001 | 0,0001 | 0,0001 |
| 0,0500 | 0,0288 | 0,0386 | 0,0462 | 0,0370 | 0,0265 | 0,0283 | 0,0274 | 0,0465 | 0,0526 | 0,0278 |  | 0,0001 | 0,0001 | 0,0001 | 0,0001 |
| 0,0973 | 0,0675 | 0,0765 | 0,0843 | 0,0192 | 0,0515 | 0,0692 | 0,0885 | 0,0930 | 0,0806 | 0,0667 | 0,0577 |  | 0,0001 | 0,0001 | 0,0001 |
| 0,0582 | 0,0336 | 0,0494 | 0,0604 | 0,0607 | 0,0424 | 0,0343 | 0,0438 | 0,0682 | 0,0782 | 0,0571 | 0,0214 | 0,0824 |  | 0,0001 | 0,0001 |
| 0,0367 | 0,0407 | 0,0415 | 0,0556 | 0,0673 | 0,0455 | 0,0348 | 0,0716 | 0,0979 | 0,1058 | 0,0642 | 0,0479 | 0,0829 | 0,0624 |  | 0,0001 |
| 0,0864 | 0,0586 | 0,0720 | 0,0882 | 0,0750 | 0,0571 | 0,0645 | 0,1042 | 0,1216 | 0,1153 | 0,0940 | 0,0661 | 0,0828 | 0,0692 | 0,0702 |  |
| 0,1195 | 0,0798 | 0,1042 | 0,0909 | 0,1047 | 0,0809 | 0,0911 | 0,1029 | 0,1177 | 0,1223 | 0,0934 | 0,0835 | 0,1149 | 0,1012 | 0,0897 | 0,0967 |
| 0,0689 | 0,0374 | 0,0509 | 0,0679 | 0,0597 | 0,0430 | 0,0420 | 0,0495 | 0,0685 | 0,0631 | 0,0480 | 0,0214 | 0,0749 | 0,0265 | 0,0571 | 0,0452 |
| 0,0581 | 0,0339 | 0,0482 | 0,0581 | 0,0344 | 0,0269 | 0,0379 | 0,0705 | 0,0891 | 0,0765 | 0,0567 | 0,0305 | 0,0525 | 0,0454 | 0,0512 | 0,0469 |
| 0,0457 | 0,0353 | 0,0490 | 0,0569 | 0,0614 | 0,0352 | 0,0350 | 0,0664 | 0,0945 | 0,0924 | 0,0650 | 0,0354 | 0,0874 | 0,0449 | 0,0435 | 0,0659 |
| 0,0567 | 0,0278 | 0,0495 | 0,0600 | 0,0471 | 0,0355 | 0,0328 | 0,0442 | 0,0604 | 0,0584 | 0,0423 | 0,0210 | 0,0681 | 0,0344 | 0,0541 | 0,0702 |
| 0,0534 | 0,0388 | 0,0546 | 0,0628 | 0,0509 | 0,0369 | 0,0404 | 0,0525 | 0,0665 | 0,0688 | 0,0443 | 0,0349 | 0,0643 | 0,0502 | 0,0440 | 0,0711 |

| SUS | WIS | GRA | UCK4 | EDS | STM |
|---|---|---|---|---|---|
| 0,0001 | 0,0001 | 0,0001 | 0,0001 | 0,0001 | 0,0001 |
| 0,0001 | 0,0001 | 0,0001 | 0,0001 | 0,0001 | 0,0001 |
| 0,0001 | 0,0001 | 0,0001 | 0,0001 | 0,0001 | 0,0001 |
| 0,0001 | 0,0001 | 0,0001 | 0,0001 | 0,0020 | 0,0001 |
| 0,0001 | 0,0001 | 0,0001 | 0,0001 | 0,0001 | 0,0001 |
| 0,0001 | 0,0001 | 0,0001 | 0,0001 | 0,0001 | 0,0001 |
| 0,0001 | 0,0003 | 0,0008 | 0,0002 | 0,0008 | 0,0016 |
| 0,0001 | 0,0003 | 0,0007 | 0,0002 | 0,0719 | 0,0001 |
| 0,0001 | 0,0001 | 0,0001 | 0,0001 | 0,0001 | 0,0001 |
| 0,0001 | 0,0153 | 0,0021 | 0,0001 | 0,0122 | 0,0011 |
| 0,0001 | 0,0001 | 0,0001 | 0,0001 | 0,0005 | 0,0001 |
| 0,0001 | 0,0001 | 0,0001 | 0,0001 | 0,0001 | 0,0001 |
| 0,0001 | 0,0001 | 0,0001 | 0,0001 | 0,0001 | 0,0001 |
| 0,0001 | 0,0001 | 0,0001 | 0,0001 | 0,0001 | 0,0001 |
| 0,0001 | 0,0001 | 0,0001 | 0,0001 | 0,0001 | 0,0001 |
| 0,0001 | 0,0001 | 0,0001 | 0,0001 | 0,0001 | 0,0001 |
| 0,0001 | 0,0001 | 0,0001 | 0,0001 | 0,0001 | 0,0001 |
| 0,0001 | 0,0001 | 0,0001 | 0,0001 | 0,0001 | 0,0001 |
| 0,0001 | 0,0001 | 0,0001 | 0,0001 | 0,0001 | 0,0001 |
| 0,0001 | 0,0001 | 0,0001 | 0,0001 | 0,0001 | 0,0001 |
| 0,0001 | 0,0001 | 0,0001 | 0,0001 | 0,0001 | 0,0001 |
| 0,0001 | 0,0001 | 0,0001 | 0,0001 | 0,0001 | 0,0001 |
| 0,0001 | 0,0001 | 0,0001 | 0,0001 | 0,0001 | 0,0001 |
| 0,0001 | 0,0001 | 0,0001 | 0,0001 | 0,0001 | 0,0001 |
| 0,0001 | 0,0001 | 0,0001 | 0,0001 | 0,0001 | 0,0001 |
| 0,0001 | 0,0001 | 0,0001 | 0,0001 | 0,0001 | 0,0001 |
| 0,0001 | 0,0001 | 0,0001 | 0,0001 | 0,0001 | 0,1688 |
| 0,0001 | 0,0001 | 0,0007 | 0,0001 | 0,0002 | 0,0001 |
| 0,0001 | 0,0001 | 0,0002 | 0,0001 | 0,0001 | 0,0001 |
| 0,0001 | 0,0001 | 0,0006 | 0,0001 | 0,0001 | 0,0003 |
| 0,0001 | 0,1120 | 0,1527 | 0,0064 | 0,0189 | 0,0001 |
| 0,0001 | 0,0001 | 0,0001 | 0,0001 | 0,0001 | 0,0001 |
| 0,0001 | 0,0001 | 0,0001 | 0,0001 | 0,0001 | 0,0001 |
| 0,0001 | 0,0001 | 0,0001 | 0,0001 | 0,0001 | 0,0001 |
| 0,0001 | 0,0001 | 0,0001 | 0,0001 | 0,0001 | 0,0001 |
| 0,0001 | 0,0001 | 0,0001 | 0,0001 | 0,0001 | 0,0001 |
| 0,0001 | 0,0001 | 0,0001 | 0,0001 | 0,0001 | 0,0001 |
| 0,0001 | 0,0001 | 0,0001 | 0,0001 | 0,0001 | 0,0001 |
| 0,0001 | 0,0001 | 0,0001 | 0,0001 | 0,0001 | 0,0001 |
| 0,0001 | 0,0001 | 0,0001 | 0,0001 | 0,0001 | 0,0001 |
| 0,0001 | 0,0001 | 0,0001 | 0,0001 | 0,0001 | 0,0001 |
| 0,0001 | 0,0001 | 0,0001 | 0,0001 | 0,0001 | 0,0001 |
| 0,0001 | 0,0001 | 0,0001 | 0,0001 | 0,0001 | 0,0001 |
| 0,0001 | 0,0001 | 0,0001 | 0,0001 | 0,0001 | 0,0001 |
| 0,0001 | 0,0001 | 0,0001 | 0,0001 | 0,0001 | 0,0001 |
| 0,0001 | 0,0001 | 0,0001 | 0,0001 | 0,0001 | 0,0001 |
| 0,0001 | 0,0001 | 0,0001 | 0,0001 | 0,0001 | 0,0001 |
|  | 0,0001 | 0,0001 | 0,0001 | 0,0001 | 0,0001 |
| 0,0820 |  | 0,0001 | 0,0001 | 0,0001 | 0,0001 |
| 0,0977 | 0,0345 |  | 0,0001 | 0,0001 | 0,0001 |
| 0,1105 | 0,0408 | 0,0349 |  | 0,0001 | 0,0001 |
| 0,0910 | 0,0264 | 0,0377 | 0,0374 |  | 0,0001 |
| 0,0775 | 0,0419 | 0,0482 | 0,0452 | 0,0323 |  |
